# Linking myelin and Epstein-Barr virus specific immune responses in multiple sclerosis: insights from integrated public T cell receptor repertoires

**DOI:** 10.1101/2024.10.23.619834

**Authors:** Sebastiaan Valkiers, Amber Dams, Maria Kuznetsova, My Ha, Ibo Janssens, Barbara Willekens, Kris Laukens, Benson Ogunjimi, Nathalie Cools, Pieter Meysman

## Abstract

The autoimmune responses in multiple sclerosis (MS), particularly those mediated by T cells targeting CNS-derived antigens, are broadly recognized. However, the defining triggers underlying these responses remain poorly understood. Epstein-Barr virus (EBV) infection has emerged as a primary risk factor for MS, suggesting a potential role for molecular mimicry in which EBV-specific immune responses cross-react with myelin antigens.

In this study, we analyzed the T cell receptor (TCR) repertoires of MS patients (n = 129) and controls (n = 94) from public datasets, to explore the relationship between EBV-specific and myelin-specific T cell responses. Our analysis identified clusters of TCRs that were significantly depleted among MS patients, many of which were associated with cytomegalovirus (CMV). By generating a library of myelin-reactive TCRs from stimulated peripheral blood mononuclear cells (PBMCs) obtained from MS patients and mapping these sequences to the public TCR repertoire database, we also uncovered a lower frequency of myelin-reactive TCRs in MS samples compared to controls in the public datasets. In addition, epitope-specificity prediction revealed a broader response to EBV-, but not CMV-derived epitopes.

Collectively, these findings underscore the complex role of chronic viral infections in MS. Particularly, they suggest that EBV-specific immune responses contribute to the dysregulation of the immune system in MS patients, potentially through mechanisms of molecular mimicry. While the broader response to EBV-derived epitopes and the lower frequency of myelin-reactive TCRs in MS samples are both associated with the disease, further research is needed to clarify the nature of this relationship. These observations suggest that viral and autoimmune mechanisms may contribute independently or interact in MS pathogenesis.

## Introduction

Multiple sclerosis (MS) is a chronic, inflammatory, demyelinating and neurodegenerative disease of the central nervous system (CNS) with a complex aetiology involving genetic, environmental, and immunological factors^1^. Despite extensive research, the precise mechanisms driving the autoimmune response in MS remain incompletely understood and are heterogeneous between patients^2^. T cells, particularly autoreactive T cells targeting myelin and other CNS-derived autoantigens, have been implicated in the pathogenesis of MS. Indeed, it has been shown that T cells from MS patients are able to recognize a range of different myelin antigens, including myelin basic protein (MBP)^3,4^, myelin proteolipid protein (PLP)^4,5^, and myelin oligodendrocyte glycoprotein (MOG)^6^. However, the primary event leading to uncontrolled autoreactive T cell responses remains unelucidated.

Evidence dating back to the 1980s has suggested a pivotal role for certain viruses, particularly Epstein-Barr virus (EBV) in MS pathogenesis^7,8^. Recent research has demonstrated that EBV infection imposes a significant (32-fold) increase in the risk for developing MS, consolidating the crucial role this virus plays in the onset of the disease^9^. Multiple theories have been proposed to explain how EBV might trigger MS. One of the leading hypotheses proposes a mechanism of molecular mimicry in which EBV-specific T cells and antibodies cross-react with self-antigens in the CNS^10–15^. One study showed that autologous co-cultures of EBV-transformed B cells and auto-proliferating (AP) T cells maintained and activated AP CD4+ T cells much more effectively, in the absence of any exogenous antigen, in HLA-DR15+ subjects with MS compared to healthy donors. In addition, RAS guanyl-releasing protein 2 (RASGRP2) was identified as a self-peptide presented by B cells that triggers the activation of CD4+ T cells^16^. Another study demonstrated that CD4+ T cells from HLA-DR15+ MS patients can respond to HLA-DR-derived self-peptides presented by B cells. These CD4+ T cells also cross-react with peptides from MBP, EBV, *Akkermansia muciniphila* (a human intestinal symbiont), and RASGRP2^17^. While the evidence for EBV and other microorganisms as a necessary trigger for the development of MS is accumulating, the exact immunopathogenic mechanisms remain elusive.

The T cell receptor (TCR) repertoire, which reflects the diversity and specificity of T cell responses, may offer valuable insights into the immune dynamics associated with EBV and myelin peptides in MS patients. Previous studies have already shown that MS patients exhibit a broader TCR repertoire against EBV peptides^18^. Moreover, TCR sequencing has successfully identified disease-associated signatures^19^ and has even led to the discovery of clinically relevant therapeutic targets in other autoimmune diseases. For example, Britanova and colleagues recently developed a highly specific immunotherapy against TRBV9+ T cells in an ankylosing spondylitis patient^20^.

In this study, we leverage integrated public TCR repertoire data to further explore the relation between EBV and a myelin-specific T cell response in MS. By analysing TCR sequences from MS patients, healthy controls, and individuals with other CNS-related pathologies, we aimed to identify MS-associated TCR signatures and assess their potential role in either increasing the risk of MS development or providing a protective effect against the disease. Our comprehensive dataset includes bulk TCR repertoire data from nine independent studies^16,21–28^, encompassing samples from peripheral blood and cerebrospinal fluid (CSF). In addition to analysing public TCR data, we generated a library of myelin-reactive TCRs by stimulating peripheral blood mononuclear cells (PBMCs) from MS patients with myelin-derived peptides.

## Methods

### Public datasets

Bulk TCRβ repertoire data was collected from 9 independent studies (Supplementary table 1). This dataset contained 827 total samples, originating from 140 MS patients, 66 healthy controls (HC) and 33 non-healthy controls (NHC). The NHC population comprised samples from patients diagnosed with non-MS CNS inflammatory disorders. This group included 14 patients with Susac syndrome (SuS), 14 with HTLV-1-Associated Myelopathy/Tropical Spastic Paraparesis (HAM/TSP), and 5 with progressive multifocal leukoencephalopathy (PML). Tissue source of the samples included peripheral blood (n=767) and CSF (n=20). The treatment regime was described for 67 out of 140 MS patients. More detailed information about the different treatments, sampling timepoints and the different datasets can be found in Supplementary table 1.

### Myelin cultures

#### PBMC Stimulation

Three subjects, diagnosed with MS, were recruited at the University Hospital Antwerp (UZA). Blood samples were taken from all three patients and frozen after collection. Ethical approval for this study was granted by the UZA ethical committee. Cryopreserved peripheral blood mononuclear cells (PBMCs) isolated from MS patients were thawed in preheated Iscove’s Modified Dulbecco’s Medium (IMDM, Life Technologies, Merelbeke, Belgium), supplemented with 10% human AB serum (hAB; Life Technologies). Next, the cell suspension was transferred into a culture flask (Greiner Bio-One, Vilvoorde, Belgium) and incubated overnight in a humidified 5% CO2 incubator at 37°C. After overnight rest, PBMCs were seeded into 12-well culture plate (Life Technologies) wells in IMDM medium supplemented with 5% hAB for six days with one of seven different myelin peptides added as stimuli. Wells with no added peptides and wells with added DMSO in a concentration equal to one in peptide-stimulated wells were used as stimulation controls. The 12-well plate layout with all the conditions per patient is shown in Supplementary figure 1.

#### Kinetics of CD4+ T cell activation

After 6 days of incubation with myelin peptides, all cells were stained with the following antibodies: anti-CD3-peridinin chlorophyll (PerCP)-Cyanine5.5 (Cy5.5) (BioLegend, Amsterdam, The Netherlands), Anti-CD4-Brilliant Violet (BV) 510 (BioLegend), Anti-CD8-Pacific Blue (PB) (BioLegend), Anti-CD71-BV786 (BioLegend), Anti-CD98-Brilliant Blue (BB) 515 (BioLegend), and viability was assessed using the LIVE/DEAD Fixable Near-IR Dead Cell Stain Kit (ThermoFisher Scientific, Merelbeke, Belgium). Next, eleven different TotalSeq™-C anti-human Hashtag oligo-antibodies (0.5 g/106cells; BioLegend, Supplementary table 2) were added to the PBMCs stimulated with myelin peptides to enable further multiplexing of the sorted samples at the single-cell TCR sequencing step. Additionally, all peptide-stimulated PBMCs were stained with TotalSeqTM-A 0390 anti-human CD127 - and TotalSeqTM-C 0085 anti-human CD25 oligo-antibodies (BioLegend, Supplementary table 3) to distinguish between conventional T cells (CD127+CD25-) and Tregs (CD127-CD25hi) in sequenced samples. Activated CD4+ T cells were sorted as CD3+CD4+ by means of Fluorescence Activated Cell Sorting (FACS) using a FACSAria II device (BD Biosciences, Erembodegem, Belgium) (Supplementary figure 2). After sorting, an aliquot of the sorted cells was taken for cell count using Countess III Automated Cell Counter and Countess™ Cell Counting Chamber Slides (Thermo Fisher Scientific). A viability of at least 90% for every sample was confirmed by trypan blue staining. Flow cytometric data were analysed using FlowJo 10.9.0 Software (Flowjo™, TreeStar Inc, Ashland, OR, USA). After FACS sorting and counting cells sorted from different peptide-stimulated conditions were pooled into one sample and placed on ice. Cell number and viability of the pooled sample were measured as described above. Then cells were centrifuged for 5 minutes at 400 x g, resuspended in phosphate-buffered saline and immediately used for the single-cell library preparation step.

### Single-cell library preparation and sequencing

Single-cell RNA sequencing libraries were prepared from the pooled samples using the 10x Next GEM 5’ Single Cell v2 Dual Index Kits with Feature Barcode technology (10x Genomics), following the manufacturer’s protocol. Specifically, Single Cell V(D)J library (containing full-length paired T-cell receptor (TCR) transcripts) and Single Cell Hashtag oligo (HTO) library (5’ Cell Surface Protein library containing TotalSeq Hashtag oligo barcodes for subsequent demultiplexing of cells labelled with different TotalSeq-C Hashtag antibodies) were prepared for each multiplexed sample.

The targeted cell recovery was up to 10,000 cells per multiplexed sample. The prepared libraries were sequenced at the VIB Nucleomics Core (Leuven, Belgium) using the NovaSeq 6000 Sequencing System (Illumina) with a sequencing depth of 5,000 read pairs per cell for each library.

### Data processing

For repertoires in PRJNA495603 and PRJNA579190, raw FASTQ files were obtained from the NCBI Sequence Read Archive (SRA). MiXCR^29^ (v4.1.0) was used to extract clonotypes from the raw sequencing data. Datasets originating from the ImmunoSEQ platform did not have the raw sequencing data available. For these repertoires, we started from the clonotype tables available in the ImmuneAccess database. Finally, we used Cell Ranger (v7.1.0) to extract paired αβTCRs from the single-cell sequencing data.

All repertoires were parsed following identical criterions to select functional TCR sequences. Any non-functional TCR whose nucleotide sequence included a stop-codon was removed. TCRs lacking a valid V/J annotation in IMGT and pseudogene or ORF V or J genes were also removed from the analysis (see https://www.imgt.org/IMGTrepertoire/LocusGenes/). In addition, TCRs with CDR3 sequences containing non-amino acid characters were excluded. Furthermore, each TCR sequence required a cysteine (C) at the N-terminal, as well as a phenylalanine (F), tryptophan (W) or cysteine (C) at the C-terminal end of the CDR3 amino acid sequence sequence. TCRs with a CDR3 containing less than 6 or exceeding 30 amino acids were excluded. For our single-cell sequencing data, any TCR sequences that were shared between ≥ 2 peptide conditions were removed in order to minimise the inclusion of non-specific or bystander activated clones.

### Diversity analysis and repertoire similarity

TCR repertoire diversity was estimated using Pielou’s evenness index^30^, calculated as:

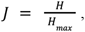

where *H*is equal to the Shannon-Wiener index.

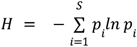

Here *S* is the total number of unique clones in the repertoire and *p*_i_ is the relative frequency of the *i*th clone. We evaluated differences in evenness across health states (MS, HC and UHC) in the PBMC samples, adjusting for age, sex and study of origin. Studies 2 and 8 were excluded from the diversity analysis due to the missing information on age and/or sex of participants. Additionally, studies 3 and 9 did not include PBMC samples and were therefore analysed separately. We used a linear mixed-effect model with health state, age, sex as fixed effects, and study of origin as a random effect. Three separate models were built for CD4, CD8, and unsorted samples to account for potential differences in TCR diversity across these subsets.

TCR repertoire similarity was measured using the Jaccard index. We used CompAIRR^31^ to calculate the pairwise Jaccard similarity between all repertoires in the public bulk TCRβ dataset.

### TCR clustering and epitope-specificity annotation

TCR repertoires from all studies were combined and TCR clustering was performed using the ClusTCR algorithm^32^. The integrated dataset excluded any of the non-PBMC samples, as well as post-treatment timepoints for the repertoires in the Amoriello 2020 cohort for patients who received AHSCT or NTZ treatment. We also excluded all the repertoires originating from samples that were collected during pregnancy in the Ramien 2019 cohort.

TCRs were annotated with epitope-specificity information. We used the ImmuneWatch (IMW) DETECT algorithm to predict epitope specificity for TCRs in the public repertoires, as well as the myelin-reactive TCRs obtained through stimulation experiments. Any predictions with a score of >0.23, as per the recommendations in the IMW DETECT documentation (https://immunewatch.gitlab.io/detect-docs/scoring), were considered reliable annotations.

### Software

All analyses were performed using Python (3.9). Bivariate statistics and hierarchical clustering was performed using the scipy (1.8) module in Python. The statsmodels (0.14) package was used for the implementation of the linear mixed effect models and multiple testing correction. For the principal component analysis (PCA), we used the implementation available in scikit-learn (1.3).

## Results

We collected data from 10 independent studies to identify MS-associated signatures that could be used for patient stratification or the development of personalised immunotherapies. Our dataset comprised a total of 791 samples: 500 samples from 141 MS patients, 220 samples from 66 healthy donors, and 71 samples from 33 patients with other CNS inflammatory disorders (Figure 1A,B). Four of the nine studies did not differentiate between the clinical subtypes of MS. In the remaining five studies, the majority of patients were diagnosed with relapsing remitting MS (RRMS), and a smaller fraction with progressive relapsing MS (PRMS), primary progressive MS (PPMS) and secondary progressive MS (SPMS). Of all the samples, 28.7% contained unsorted T cells or unspecified subtypes. CD4+ samples accounted for 34.4%, with 5.2% sorted into the naive fraction and 8.3% identified as CD4+ memory T cells. Within the CD4+ memory category, 2.4% were central memory (CM), 2.3% were effector memory (EM), and 3.9% were unspecified memory T cells. The remaining 36.9% of the samples contained CD8+ T cells. Among these, 5.6% were naive, 8.6% were memory T cells (including 2.4% CM and 2.3% EM), and 2.4% were effector memory cells re-expressing CD45RA (T_EMRA_) cells (Figure 1B).

**Figure 1.**
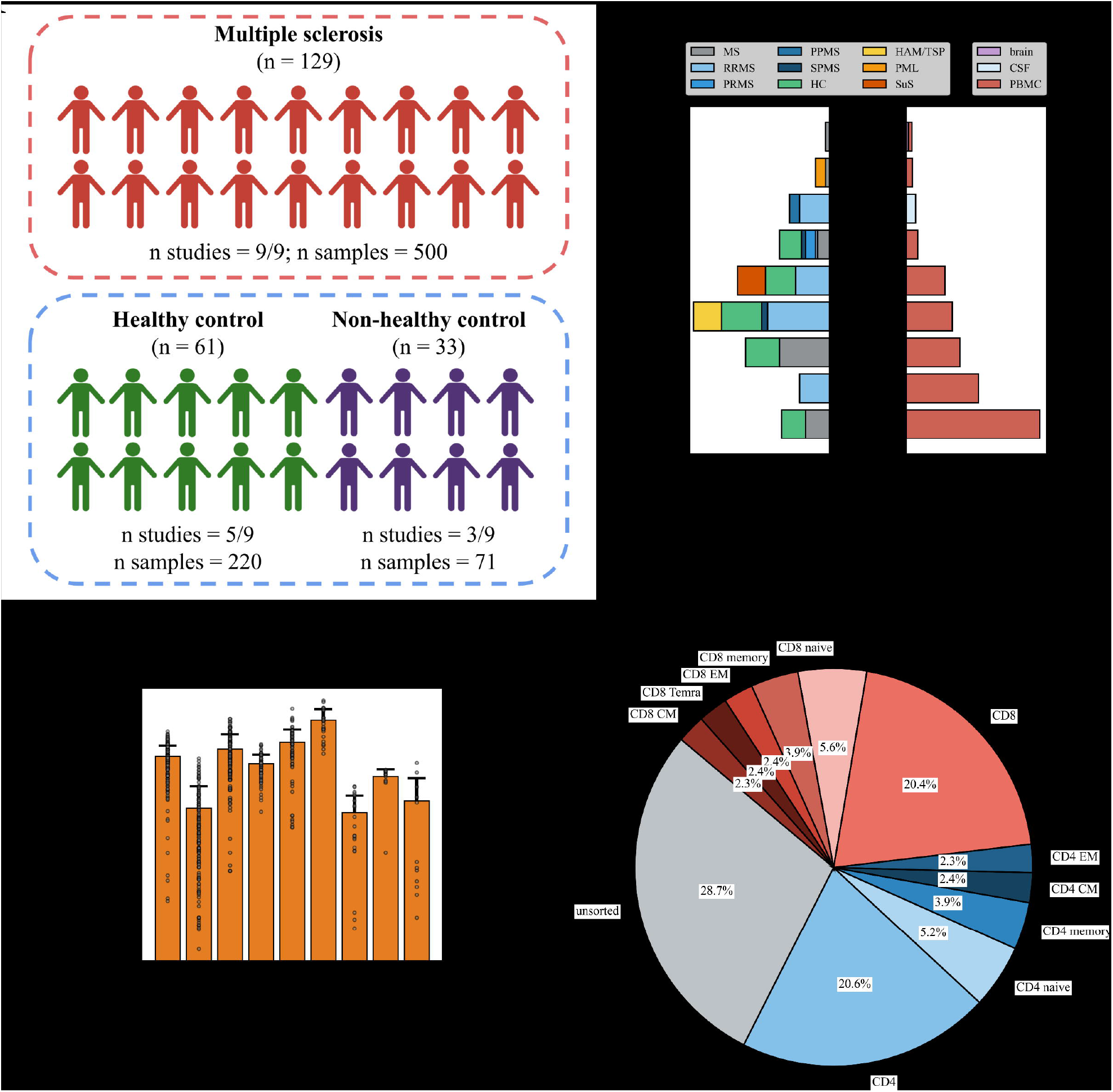
Public TCR repertoire data from MS studies. **A**. Overview of the complete public dataset of MS, HC and NHC subjects. **B**. Number of subjects and samples across different studies, categorised by diagnoses and tissue sources. The left panel represents the number of patients per study, with bars stacked to differentiate between various diagnoses. The right panel illustrates the total number of samples per study, with bars stacked according to different tissue sources. **C**. Average number (+ 1 std. dev) of VDJ rearrangements per study. **D**. Different T cell subsets across all studies.

The number of TCR rearrangements varied strongly between datasets, which is likely the result of differences in sequencing depth, tissue source (blood, CSF), and whether or not the samples were sorted into various T cell subsets. For instance, the low number of rearrangements per sample in the Amoriello 2021 dataset resulted from the samples originating from the CSF and being sorted into different subsets: CD4 CM, CD4 EM, CD4 naive, CD8 CM, CD8 EM, CD8 naive, and CD8 T_EMRA_. In contrast, the samples in the Shugay 2015 dataset are unsorted PBMCs. Consequently, the latter contained approximately a 100-fold more unique TCR sequences on average per repertoire compared to the former (Figure 1B). Overall, the MS samples in our dataset contained fewer rearrangements than both control groups (Supplementary figure 3A). Lastly, we also evaluated the age distribution of the subjects to ensure any differences found between the groups are not a direct result of age-related factors such as accumulation of chronic virus-specific TCRs or T cell exhaustion. The ages of the participants across the three different groups (MS, HC, NHC) were distributed differently (p=1.67e-05, one-way ANOVA). While there was no significant difference in age between the MS and HC groups (p=0.24, t-test), the NHC participants tended to be slightly older than the MS participants (p=0.034, t-test). Expectedly, we also observed an age-related decline in the number of unique TCRs per repertoire. However, the rate of decline was indifferent between groups (Supplementary figure 3B & Supplementary table 4).

### Biased V gene usage in public data set is associated with study and cell type

To study the V gene usage in the bulk data set, we performed PCA on the V gene frequencies in each repertoire prior to V gene parsing. There was a clear distinction in the V gene usage pattern among repertoires that were surveyed using the ImmunoSEQ assay, compared to 5’RACE protocols (Figure 2A). Particularly TRBV5-4, TRBV12-5 and TRBV19 were more prominent in ImmunoSEQ repertoires, while TRBV20-1 and TRBV5-1 seemed to have a stronger association to 5’RACE-based repertoires (Figure 2B). Interestingly, PCA was able to separate CD4+ and CD8+ subsets in all datasets (Supplementary figure 4). This separation was driven by a selection of V genes. More specifically, TRBV20-1, TRBV7-2 and TRBV5-1 are negatively correlated with CD8+ samples, while TRBV27, TRBV7-9 and TRBV4-1 tended to have a positive association with the CD8+ phenotype. The unsorted samples were typically more CD4-like in terms of V gene usage (Supplementary figure 4), consistent with the evidence that CD4+ T cells typically represent a larger fraction of the total T cell population compared to CD8+ T cells^33^. In addition to the differences observed between CD4+ and CD8+ subsets, we identified differential V gene usage between MS patients and healthy controls in the Emerson datasets (Supplementary figure 4,5). However, closer examination of the T cell subset annotations suggests that this separation is likely due to a batch effect. PCA, with PC1 accounting for approximately 57% of the variance, further separated regular CD4+/CD8+ samples from those with naive and memory subtypes, resulting in four distinct clusters: CD4+, CD4+ naive/memory, CD8+, and CD8+ naive/memory (Supplementary figure 4). Collectively, these observations highlight the importance of considering both technical and biological variation when interpreting differences in V gene usage between samples.

**Figure 2.**
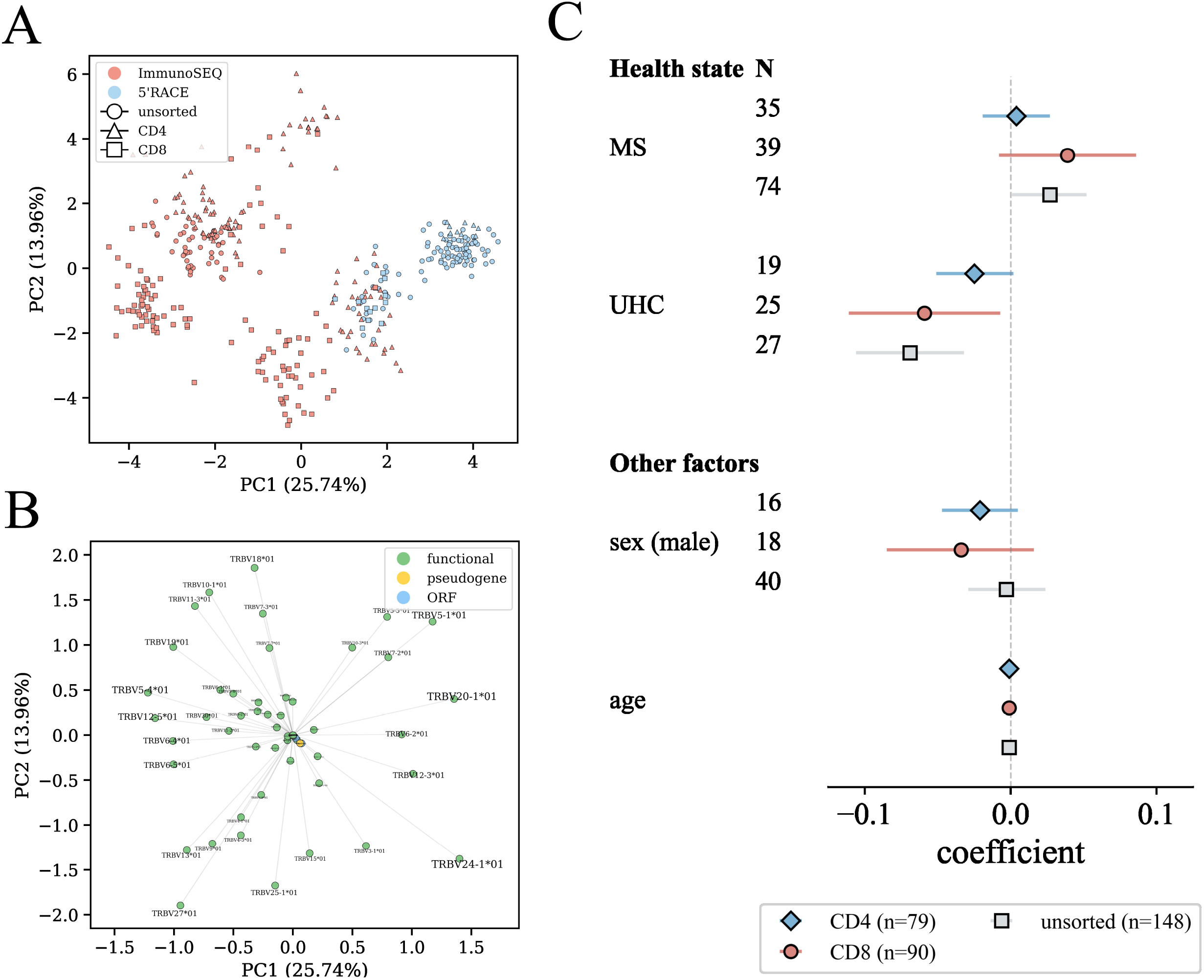
Analysis of public TCR repertoires from MS patients and controls. **A**. PCA based on V gene frequency across 6/9 datasets. The PCA plot was annotated by T cell fraction and library preparation technology per sample. There is a clear difference in V gene usage between samples prepared using ImmunoSEQ versus 5’RACE-based protocols. **B**. Loadings plot indicating the contribution of each V gene to the principal components. **C**. A generalized linear mixed effect model was used to model TCR repertoire diversity between HC, NHC and MS patients, including age and sex as fixed, and study as random effects. A model was built for CD4+, CD8+ and unsorted T cells separately. No differences in TCR repertoire diversity were found between MS patients and HCs. A slight increase in diversity was found among unsorted T cells in MS patients (compared to HCs). CD8+ and unsorted T cells in NHCs were found to have lower diversity compared to HCs. No significant effects were observed for sex or age.

### TCR repertoire diversity and overlap in MS is not dissimilar to healthy controls

We next analysed TCR repertoire diversity between HC, MS and UHC (Figure 2C). The analysis was conducted separately on CD4+, CD8+, and unsorted T cells. We found no difference in repertoire diversity for CD4+ and CD8+ T cells between MS patients and HC. However, in unsorted PBMC samples, diversity was slightly higher in MS patients compared to controls (95% CI = [0.001, 0.052]). In contrast, we found lower diversity for non-healthy controls relative to healthy subjects in the unsorted TCRs (95% CI = [-0.106,-0.032]) as well as the CD8 fraction (95% CI = [-0.111,-0.007]). While we observed an age-related decline in species richness (Supplementary figure 3B), there was no significant effect for age on repertoire diversity in this selection of datasets when comparing differences between subjects with different health states. Indeed, when evaluating species richness individually in each group, we see a similar age-related decline in MS, HC and NHC (Supplementary table 4). We next compared the pairwise Jaccard similarity between all repertoires in the public dataset using CompAIRR, in order to identify potentially differentiating patterns between MS and controls. We found that repertoires clustered largely by study, and not by health state (MS, HC, NHC) or tissue type (Supplementary figure 6).

### TCR clustering identifies MS depletion-associated signatures

TCR clustering was performed using ClusTCR on a combination of all repertoires in the bulk TCRβ dataset (37,346,317 total TCRs < 21,119,263 MS, 11,825,847 HC, 4,401,207 NHC). We identified the presence or absence of each cluster in every repertoire and performed an enrichment test to evaluate the association or depletion of clusters between MS patients and controls. None of the MS-associated clusters reached the threshold for significance (Figure 3A). However, we identified 166 clusters that were significantly depleted in MS patients. There was a distinct pattern in TRBV usage between MS-depleted and MS-associated clusters (Figure 3B). Upon inspecting the V gene usage of the top 100 depleted clusters, we found a predominant enrichment for TRBV20, and to a lesser extent also TRBV6, 7, 5, 24, and 11. In contrast, the top 100 MS-associated clusters primarily included TRBV6 and 30, with smaller amounts of TRBV11, 2, and 25. Moreover, many of the top 100 significantly depleted clusters were very large (> 500 sequences), with high P_gen_ sequences indicating common rearrangements (Supplementary figure 7A,B). We next matched the sequences from the MS-depleted clusters with the recently published CMV Exposure Co-Occurrence (ECO) cluster to infer CMV association and HLA-restriction. We found that 44 out of the 166 significantly depleted clusters had at least 1 member that matched with TCR clones in the CMV ECOcluster.

**Figure 3.**
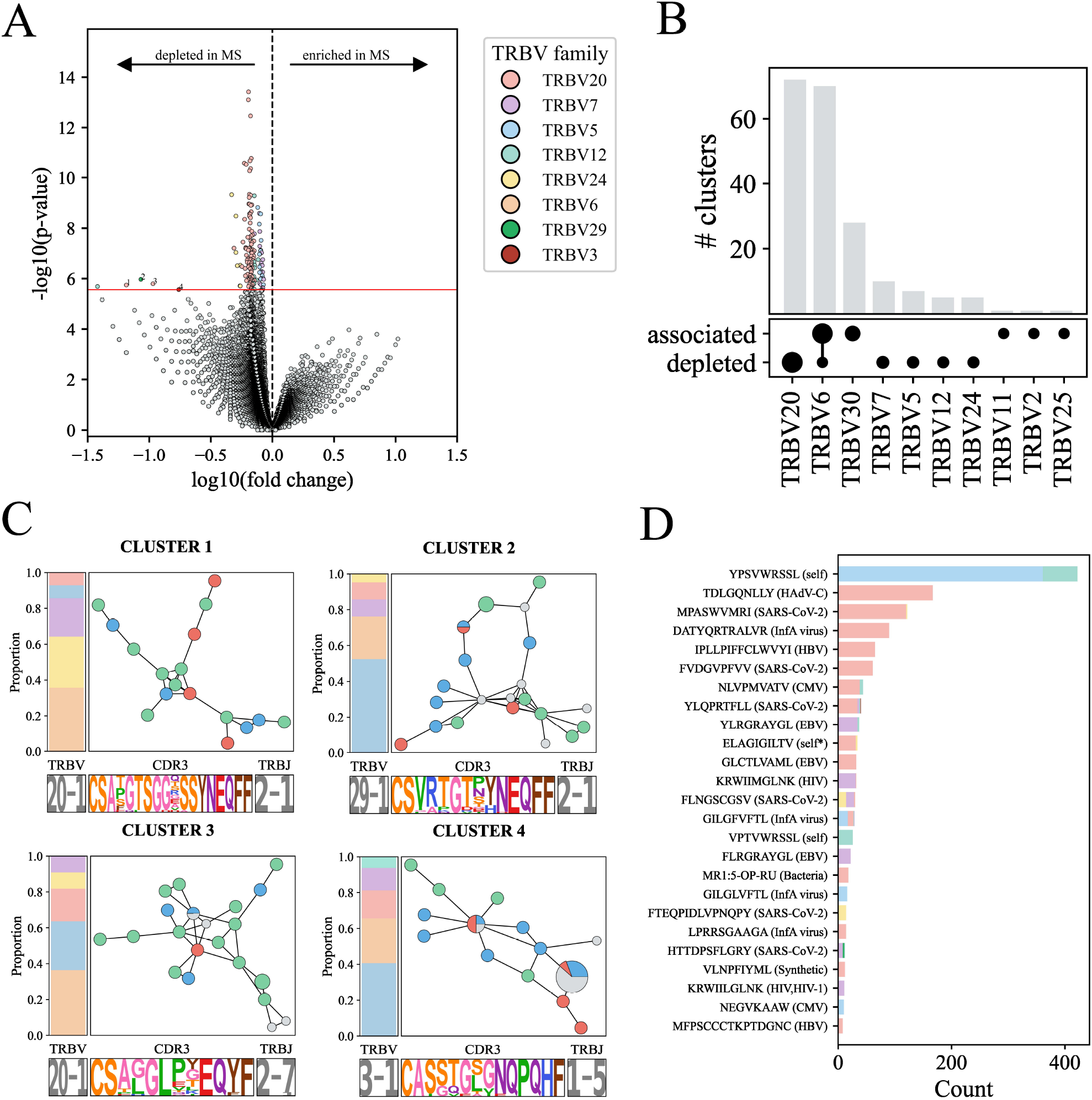
TRB clusters association analysis between MS patients and HC. **A**. Volcano plot showing the fold enrichment of the TCR cluster in MS patients versus the Fisher’s p-value for the association. The red line indicates the selected p-value threshold based multiple testing correction (Benjamini-Hochberg procedure). Significant clusters were coloured by TRBV family. None of the clusters reached significant enrichment in MS patients. In contrast, 166 clusters were significantly depleted in MS. 4 clusters were selected for further inspection (in C.), based on p-value, fold enrichment, and presence across multiple datasets. **B**. Comparison of the 166 significant clusters with the 166 most MS-enriched clusters (by p-value). The clusters most associated with MS primarily contained TRBV6 and TRBV30, and to a lesser extent also TRBV11, TRBV12 and TRBV25. Clusters depleted in MS patients primarily had TRBV20, and smaller amounts of TRBV6, TRBV7, TRBV12 and TRBV14. The grey nodes indicate that this TCR was present in one of the NHC samples, but not in MS or HC. **C**. Annotated clusters 1-4 in from A. **D**. Epitope-specificity annotation of all TCRs in the 166 significantly depleted clusters using the IMW DETECT model. * = synthetic human peptide.

Among the 166 clusters, 5 showed > 5-fold enrichment in control subjects compared to MS patients (Figure 3C). The TRBV12-3 cluster contained only 3 sequences, and was exclusive to the Sousa dataset. However, the other 4 remaining clusters were each shared among 5 different datasets (4 annotated clusters in Figure 3A). The clusters had the following characteristics (V_CDR3_J): TRBV20-1_CSAXGTSGGXSSYNEQFF_TRBJ2-1, TRBV29-1_CSVRTGTX[YH]NEQFF_TRBJ2-1, TRBV20-1_CSA[GL]PXEQYF_TRBJ2-7, TRBV3-1_CASSTG[SL]GNQPQHF_TRBJ1-5. All of the clusters originated from a mix of CD4+ and CD8+ repertoires. Next, we used IMW DETECT in an attempt to reveal the epitope specificity for the clusters but found no reliable annotations for any of the TCRs in clusters 1-4. Moreover, none of these clusters were found to match sequences in the CMV ECOcluster.

When running IMW DETECT on the entire set of TCRs in the 166 MS-depleted clusters, we observed the strongest annotation for the human YPSVWRSSL peptide, a mutant of a cancer neo-epitope derived from RNF43 (Figure 3D). Upon further inspection, this recognition was mainly driven by one cluster of TRBV12 and one of TRBV5-4 TCRs. When looking at the total number of unique clusters (rather than unique TCR sequences) recognizing each epitope, we identified the highest annotation for CMV epitopes ELAGIGILTV (6.6% of clusters), NLVPMVATV (5.4% of clusters), and the SARS-CoV-2 epitope MPASWVMRI (5.4% of clusters) (Supplementary figure 7C). Next, we were interested to see if any of the TCRs in the significantly MS-depleted clusters would provide information towards treatment outcome. For this, we looked at the number of approximate (TCRdist ≤ 12) hits before and 24 months after treatment of 7 MS patients with AHSCT (Amoriello 2020 cohort). Interestingly, we saw a 10-fold increase in depletion-associated TCRs after treatment (pre: 49/514,954 clones; post: 423/421,762; p=8.02e-93, Fisher’s test). The increase in the post-treatment samples was most prominent in patient MS032 (Supplementary figure 7D). We then zoomed in at the dynamics of the subset of strongly depleted clusters (1-4) in this patient population. Again, this increase was mainly observed in subject MS032, mainly consisting of TCRs from the TRBV29-1 cluster. Interestingly this cluster predominantly exhibited a CD8+ T cell memory phenotype at 24 month post-treatment in MS032, while primarily CD4+ T cells in other patients. It is important to note, however, that MS032 experienced a relapse during the 2 year follow-up period.

### Identification of myelin-reactive TCRs in MS patients

Little is known about the myelin-reactive repertoire in MS patients. In order to gain insight into the myelin-reactive properties of the TCR repertoire in MS patients, we extracted PBMCs from MS patients and stimulated them with 7 myelin peptides (MBP1-4, MOG1-2, PLP1). About half of the cultures had activity higher than controls (Supplementary figure 8,9). We next determined the TCRαβ of CD154+ T cells using the Chromium™ Single Cell Immune Profiling Solution by 10X Genomics. After filtering, we were left with 807 unique clones with productive αβ pairs (Figure 4A).

First, we looked at the sequence similarity among the sequences in order to identify any clusters. We identified large heterogeneity between the sequences, even within the same peptide pool (Supplementary figure XX). In fact, when clustering the sequences based on their pairwise TCRdist distances, TCRs originating from the same peptide pools did not seem to group together (Supplementary figure 10). Again, we used the IMW DETECT method to predict epitope specificity of the myelin-reactive TCRs. Surprisingly, there was a substantial proportion (24/807) of potential mucosal-associated invariant T (MAIT) cells expressing TRAV1-2/TRAJ33 that also had predicted specificity for MR1:5-OP-RU. Any clones with MAIT characteristics were removed from the dataset.

**Figure 4.**
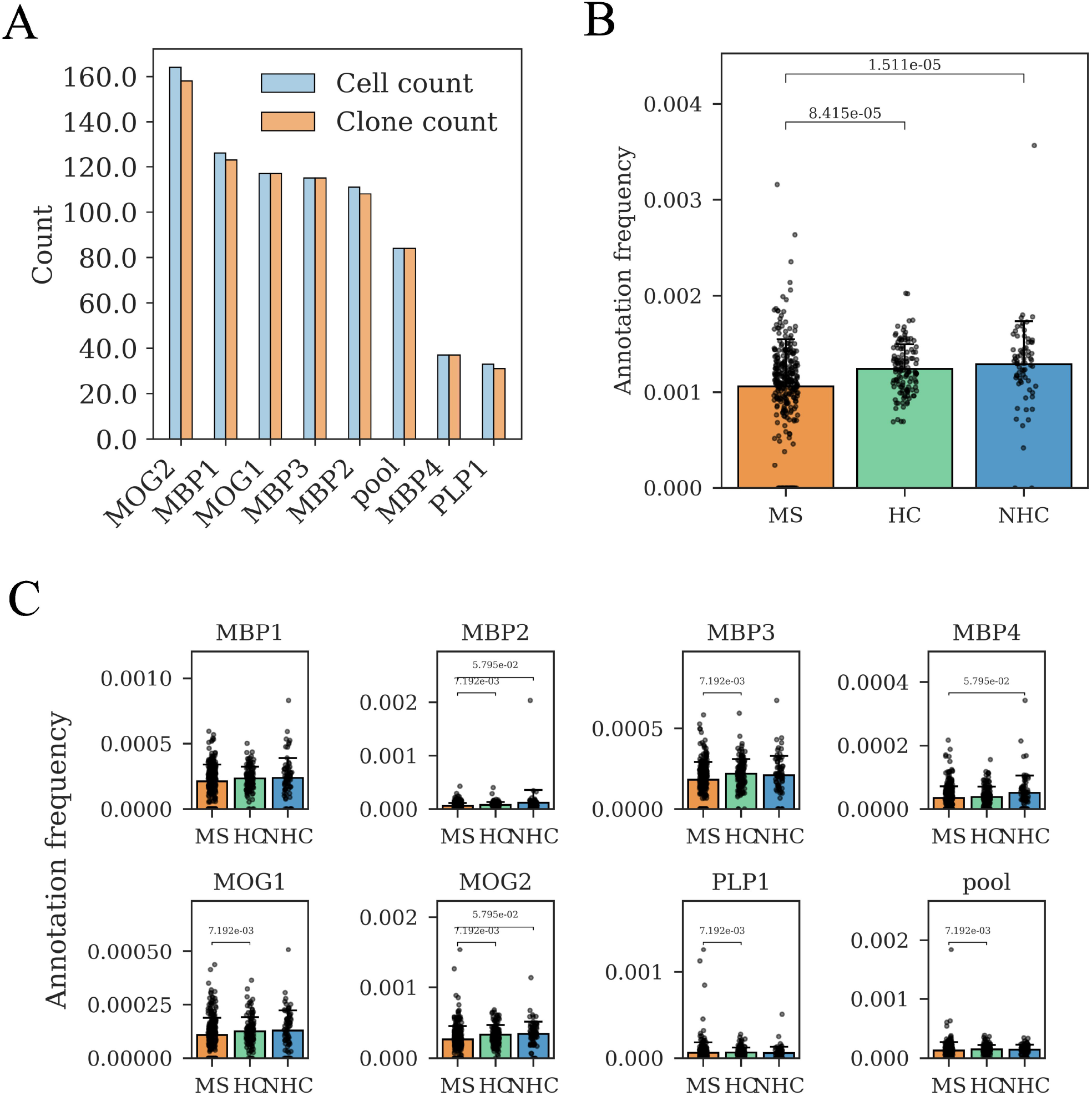
Identification of myelin-reactive TCRs in repertoires of MS, HC and UHC subjects. **A**. Total number of identified cells and unique T cell clones reactive for different myelin peptides. **B**. Annotation of myelin-reactive TCRβ sequences against a dataset of repertoires from MS, HC and NHC subjects.An approximate matching approach (TCRdist < 12.5.) was used to match TCRβ in the myelin database against sequences in the bulk dataset. Fewer myelin-reactive TCRβs were found in the MS group compared to healthy controls (p=8.415e-05, MWU test) and disease controls (p=1.511e-05, MWU test). **C**. Annotation of TCR repertoires from MS, HC and UHC subjects with TCRs reactive to myelin-derived peptides MBP1-4, MOG1-2 and PLP1. We found a significant difference in annotation frequency for MBP2-, MBP3-, MOG1-, MOG2-, and PLP1-reactive TCRs between MS and HC samples. The pooled TCRs in the pooled epitope also had significantly fewer annotation frequency in MS samples compared to HC.

We were then interested to see if any of the TCRβ chains in our myelin-reactive TCR dataset overlapped with the CD4+ myelin-specific TCRs from subject MS15 in the Ramien cohort. However, we did not observe any overlap among the two sets.

Next, we were interested to see if we could identify more myelin-specific TCRs among MS patients versus controls by matching our set of myelin-reactive TCRs obtained from MS patients to all repertoires in the public dataset. We found no difference in annotation between controls and MS patients when using an exact matching strategy (Supplementary figure 11A). However, when allowing a TCRdist ≤ 12 mismatch between the myelin-reactive TCR sequences and the TCRs in the public database, we found lower rates of annotation in MS samples compared to samples from healthy controls (p=8.415e-05, MWU test) and non-healthy controls (p=1.511e-05, MWU test) (Figure 4B). This result was indifferent between CD4+ and CD8+ fractions (Supplementary figure 11B). No differences were found between blood and CSF in terms of myelin-reactive TCR frequency in the repertoire (Supplementary figure 11C). To find out which specific epitopes were driving the differences in annotation rate between the groups we performed the annotation separately for each peptide-reactive pool of TCRs. The results of this analysis showed that, for all but two epitopes (MBP1 and MBP4), a higher proportion of approximate matches were found among HC compared to MS subjects. Additionally, for the MBP2, MBP4 and MOG2 epitope-reactive clones we found significantly more matches among the NHC cohort compared to the MS subjects (Figure 4C).

Finally, we used the same annotation strategy with the myelin-specific TCRs that were obtained from activated CD4+ T cells in the Ramien cohort. Again, no differences were observed among the groups when using an exact matching strategy (Supplementary figure 12A). Although not statistically significant, approximate matching showed lower numbers of myelin-specific TCRs among MS patients (Supplementary figure 12B), consistent with our own dataset.

### CD8+ T cells in MS patients are enriched for EBV-specific TCRs

Viral infections, especially those caused by EBV, have been proposed as one of the most plausible triggers for the onset of MS pathogenesis. Consequently, our objective was to determine whether there is an enrichment of virus-specific TCRs in the repertoires of MS patients compared to controls.

To achieve this, we utilised the IMW DETECT algorithm to predict epitope specificity across all TCR repertoires in our public database. We confidently assigned epitope labels to a total of 156,599 sequences. The highest number of sequences (27,822) were specific for the HLA-A*0201 Influenza A virus epitope GILGFVFTL. Additionally, we identified high numbers of TCRs specific for SARS-CoV-2 epitopes, particularly YLQPRTFLL, FLNGSCGSV, and (H)TTDPSFLGRY.

Next, we analysed the differences in the number of viral-specific TCRs between MS samples and those from a control population, focusing primarily on the epitope-specific responses against CMV and EBV. As a control, we also examined HIV-specific TCRs, under the assumption that the rates of HIV infection were negligible or equal among both MS and control subjects. We found that samples originating from MS patients contained a higher frequency of EBV-(p=0.0046, ranksum test), but not CMV-(p=0.091) or HIV-specific (p=0.59) TCRs compared to control samples (Figure 5A). Expectedly, these responses were primarily CD8-mediated (Figure 5B). Indeed, comparing the EBV-specific response for CD4+ and CD8+ TCR repertoires separately revealed significantly higher annotation rates in MS for CD8-associated TCRs (p=0.0095), but not among CD4+ T cells (p=0.51). We also tested PBMC and CSF samples but found no significant differences in EBV-specific TCR frequencies between the two tissue types (Figure 5C). Moreover, we observed the same predicted specificity pattern within the CSF samples as compared to the PBMCs, with most TCRs being specific to GILGFVFTL (Figure 5D).

**Figure 5.**
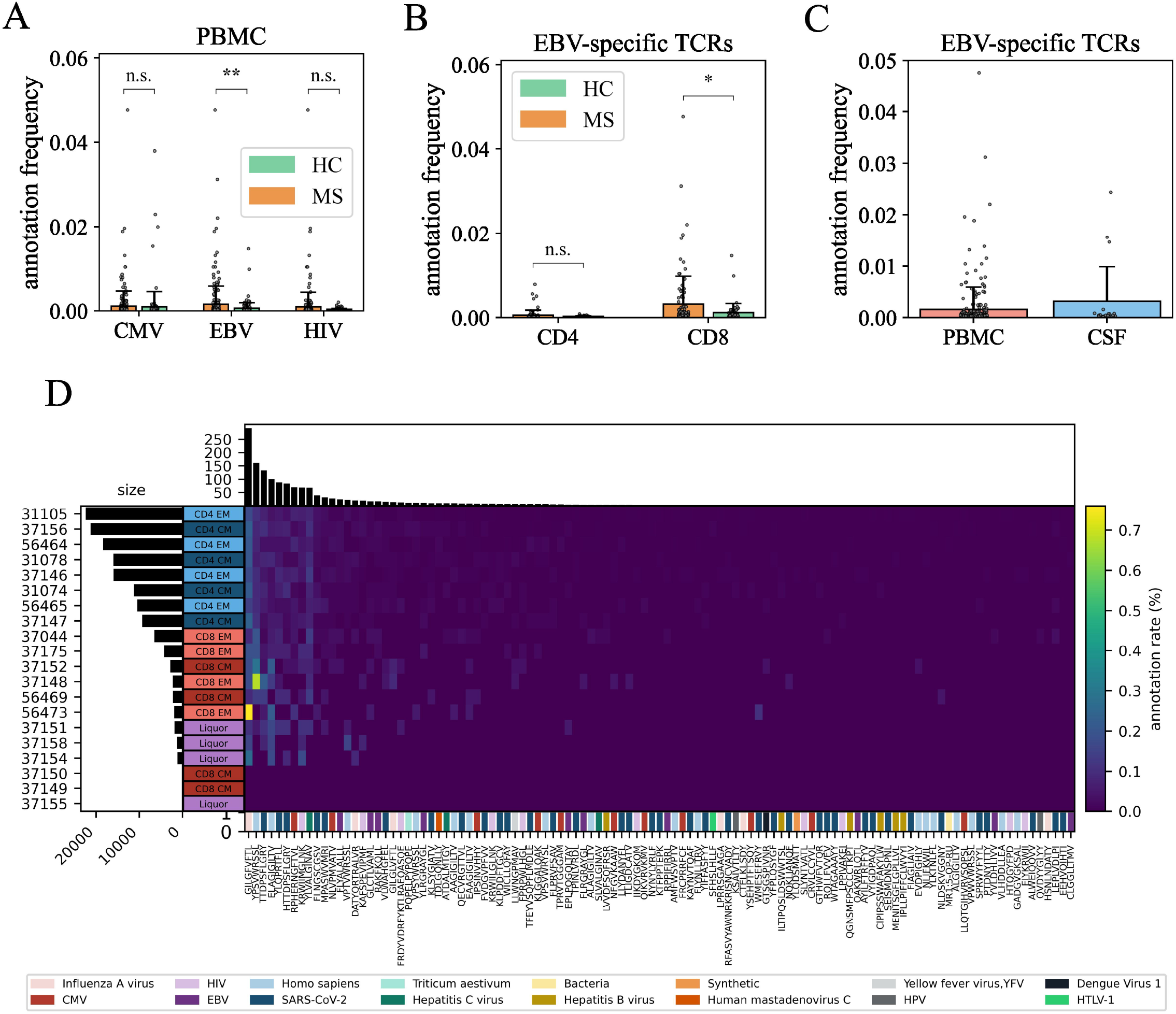
Prediction of epitope specificity in public TCR repertoires using IMW DETECT. **A**. TCR-epitope prediction in PBMC samples for MS patients and controls (HC and NHC) for CMV, EBV and HIV epitopes. The latter virus was used as a control. The bar chart shows the average annotation frequency and standard deviation. No differences were observed in overall specificity against CMV or HIV. MS patient repertoires had elevated specificity against EBV epitopes (p=0.0046, ranksum). **B**. Comparison of EBV specificity in CD4+ and CD8+ repertoires between MS and control samples. A significant increase in EBV specificity is observed in the CD8+ repertoires (p=0.0095), while no significant difference is noted in the CD4+ repertoires (p=0.51). **C**. Comparison of EBV specificity annotation rates between PBMC and CSF samples. Although CSF repertoires exhibited higher annotation rates than PBMC repertoires, the difference was not statistically significant (p=0.90). **D**. Epitope specificity heatmap for CSF repertoires from MS patients.

## Discussion

MS is a complex autoimmune disease of the central nervous system (CNS), characterised by the breakdown of myelin sheaths around nerve fibers. T cells have long been implicated in the aetiology of MS, playing a pivotal role in the autoimmune response that leads to demyelination. The TCR repertoire can provide crucial insight into the different mechanisms by which our immune system binds and neutralises target antigens. However, TCR sequencing studies in MS patients remain small-scaled and survey only a fraction of the information present in the data. Here we leveraged the public TCR repertoire in MS to uncover important signatures that may help us better understand the disease. We collected TCR repertoire data from nine independent studies, with the aim of discovering disease-associated repertoire signatures that could help us devise novel and personalised therapies for controlling this disease. Reports on the TCR diversity in MS have been highly inconsistent, with some studies suggesting lower, and others higher TCR repertoire diversity in patients with the disease. By integrating data from six different studies, we found no significant differences in repertoire diversity between MS patients and healthy controls, while the non-healthy control subjects showed decreased repertoire diversity. These findings further support the hypothesis that MS may be driven by specific autoreactive T cells rather than a general change in overall TCR diversity. However, this analysis is far from conclusive. For example, data on the sampling timepoints in these studies is extremely sparse so it is unclear in which phase of the disease the sample was taken (e.g. during relapsing or remitting phase). Moreover, studying the V gene usage in the public dataset, we identified several important biases related to technology and cell type. While we corrected for the latter in our repertoire diversity model, the former may pose a significant source of bias in comparing TCR repertoire diversity between the different groups.

TCR clustering analysis on the integrated dataset revealed no significantly MS-associated clusters. However, we did identify 166 clusters that were significantly depleted in MS samples. Interestingly, ∼27% of significantly depleted clusters in MS were found to contain at least 1 CMV-associated TCR. There is accumulating epidemiological evidence that links CMV seropositivity with reduced risks for developing MS^34–36^. While the exact mechanism for this remains elusive, this may partly explain the large number of potentially CMV-associated clusters among controls versus MS patients. Although the majority of the MS-depleted clusters were likely to be a result of high recombination probabilities or a direct effect of HLA restriction, we identified four distinct TCR clusters that showed consistent > 5-fold enrichment in control subjects in multiple studies. We hypothesised that these clusters may be primarily characterised by T_reg_ cells. However, upon closer inspection, they contained a mix of CD4+ and CD8+ T cells. We found a 10-fold increase in the frequency of these clusters in MS patients 24 months after AHSCT compared to their pre-therapy repertoires. The steep increase in the frequency of these TCRs was primarily observed in a single individual who unfortunately experienced one relapse during the post-treatment follow-up period. However, the TCRs in this individual primarily had a cytotoxic profile, compared to a CD4+ T helper phenotype in other patients. While these results remain largely inconclusive, we may speculate that the enriched TCR clusters in control subjects and their re-emergence post-AHSCT may play a role in immune surveillance and maintaining immune homeostasis. Their depletion in MS patients might contribute to the lack of effective immune regulation and control, leading to disease progression. The presence of the identified TCR clusters as primarily CD8+ T cells in the relapsing patient, versus CD4+ T cells in other patients, suggests that the function of these T cell subsets plays a crucial role in disease progression and immune response. Specifically, this differential phenotype indicates that CD8+ T cells may have a more active cytotoxic role in mediating relapse, while CD4+ T cells in other patients may exert a more regulatory or suppressive effect, contributing to disease control and remission.

We next attempted to generate a library of TCRs from activated T cells upon stimulation with myelin-derived peptides. Mapping these TCRs to bulk repertoires we did not observe an increased frequency of myelin-reactive TCRs in MS patients and we also saw no overlap with the clusters that were depleted in MS patients. This finding aligns with previous studies suggesting that the mere presence of myelin-reactive T cells is not necessarily higher in MS patients but that their functional properties and regulatory environment may be more critical factors in disease progression^37,38^. We can hypothesise that the increase in myelin-reactive clones among control subjects may indicate a more robust regulatory response against these peptides. Additionally, alternative CNS proteins such as GlialCAM^11,39^, alpha B-crystallin^14^, and others for which we did not evaluate the T cell response, could also play a role.

The role for EBV-specific T cells has been unclear, with some studies even suggesting lower, dysregulated T cell responses against EBV peptides in MS patients^40,41^. Other studies have explored the EBV-specific response through the lens of the TCR repertoire and found that MS patients exhibit a broader EBV-specific TCR repertoire compared to controls, particularly among MHC class I-restricted TCRβ sequences^18^. Indeed, our study confirms this observation by showcasing that the frequency of predicted EBV-specific, but not CMV-specific, TCRβ sequences derived from CD8+ T cells was altered in MS patients versus HCs. More specifically, MS patients had a higher frequency of TCRs specific to the Latent membrane protein 2A (LMP2A) derived epitope FLYALALLL. Interestingly, a phase I clinical trial with autologous EBV-specific T cell therapy targeting EBV nuclear antigen 1 (EBNA1), latent membrane protein 1 (LMP1), and LMP2A was shown to be a safe and effective approach for treating progressive MS, with significant clinical improvements observed in patients with strong EBV-specific T cell reactivity^42^. Furthermore, we identified no difference in the frequency of EBV-specific TCRs between CSF and PBMC in MS patient samples, suggesting that these T cells are distributed similarly across both compartments. Importantly, this highlights that EBV-specific TCRs are indeed present in the CSF, underscoring a crucial observation that may lend support to the molecular mimicry hypothesis in MS. The presence of these TCRs in the CSF suggests that the immune response to EBV is not confined to peripheral blood but extends into the CNS. However, it is important to note that our study did not include CSF data from HCs, limiting our ability to draw definitive conclusions about the uniqueness of this finding to MS patients.

Finally, we would like to highlight several limitations associated with this study. First of all, the relatively small size of our overall dataset, combined with significant batch effects among samples from various studies, may have restricted our ability to identify any strong MS-associated TCR signatures. Moreover, the public dataset suffered from a class imbalance between MS patients and (healthy) controls. This may have influenced the statistics, particularly in cluster enrichment. For example, the total number of samples used for this analysis was 451, of which only 130 originated from healthy controls. Future research with larger cohorts and better stratification of patients by clinical subtypes, as well as more detailed information about treatment histories will be essential to validate these findings and uncover more precise TCR signatures associated with MS. Next, we also detected several signs of non-specific activation in our myelin peptide cultures. Flow cytometry revealed elevated levels of T cell activation in approximately half of the cultures compared to the control group. Among these cultures with increased activation, the rise above control levels was modest. Secondly, previous research has clearly established that TCRs in epitope-specific repertoires exhibit higher levels of sequence similarity compared to non-specific repertoires^43^. Based on this, we expected to find clusters with high sequence similarity in our myelin peptide-reactive TCR pools. However, we did not observe such clusters. Thirdly, we observed several TCRs with MAIT characteristics (TRAV1-2+TRAJ33/12/20) in our myelin-reactive dataset. Indeed, epitope-specificity annotation revealed annotation with MR1:5-OP-RU for these clones. Collectively, these results highlight the need for robust validation of the activation status of the cultured T cells. This may be achieved through restimulation, such as done in Jelcic et al. (2018)^16^. Alternatively, single-cell transcriptomics may be used to differentiate between conventionally activated T cells and bystander activation through quantification of activation markers including CD69, CD25 (IL2RA), and CD40L, along with specific cytokines such as IFN-γ, IL-2, and TNF-α. Conversely, one may look to identify molecular factors that induce bystander activation such as type I IFNs, IL-12, IL-15 and IL-18^44,45^.

In conclusion, our study has revealed several critical insights into the TCR repertoire in MS. Notably, we identified a lack of significant differences in overall TCR repertoire diversity between MS patients and healthy controls, supporting the idea that the pathology of MS may be influenced by the functional profiles of specific autoreactive T cells rather than broad immune dysfunction^27,46^. In relation to this, we showed that TCR repertoires of MS patients contained significantly lower frequencies of myelin-reactive TCRβ sequences. Conversely, we also observed a unique profile in relation to EBV, with MS patients exhibiting broader EBV-specificity in the TCR repertoire compared to controls, potentially providing support for a molecular mimicry hypothesis in MS pathogenesis. Collectively, these findings illustrate the dual role of viral and autoimmune factors in MS, where the depletion of protective TCR clusters and the reduced prevalence of myelin-reactive TCRs point towards specific disruptions in the immune landscape. We propose that future research should focus on uncovering the functional properties of these dysregulated T cell populations in MS and explore how they can be targeted to restore immune homeostasis and prevent disease progression.

