## Supplementary figures and images for "Linking myelin and Epstein-Barr virus specific immune responses in multiple sclerosis: insights from integrated public T cell receptor repertoires"

### Supplementary figure 1

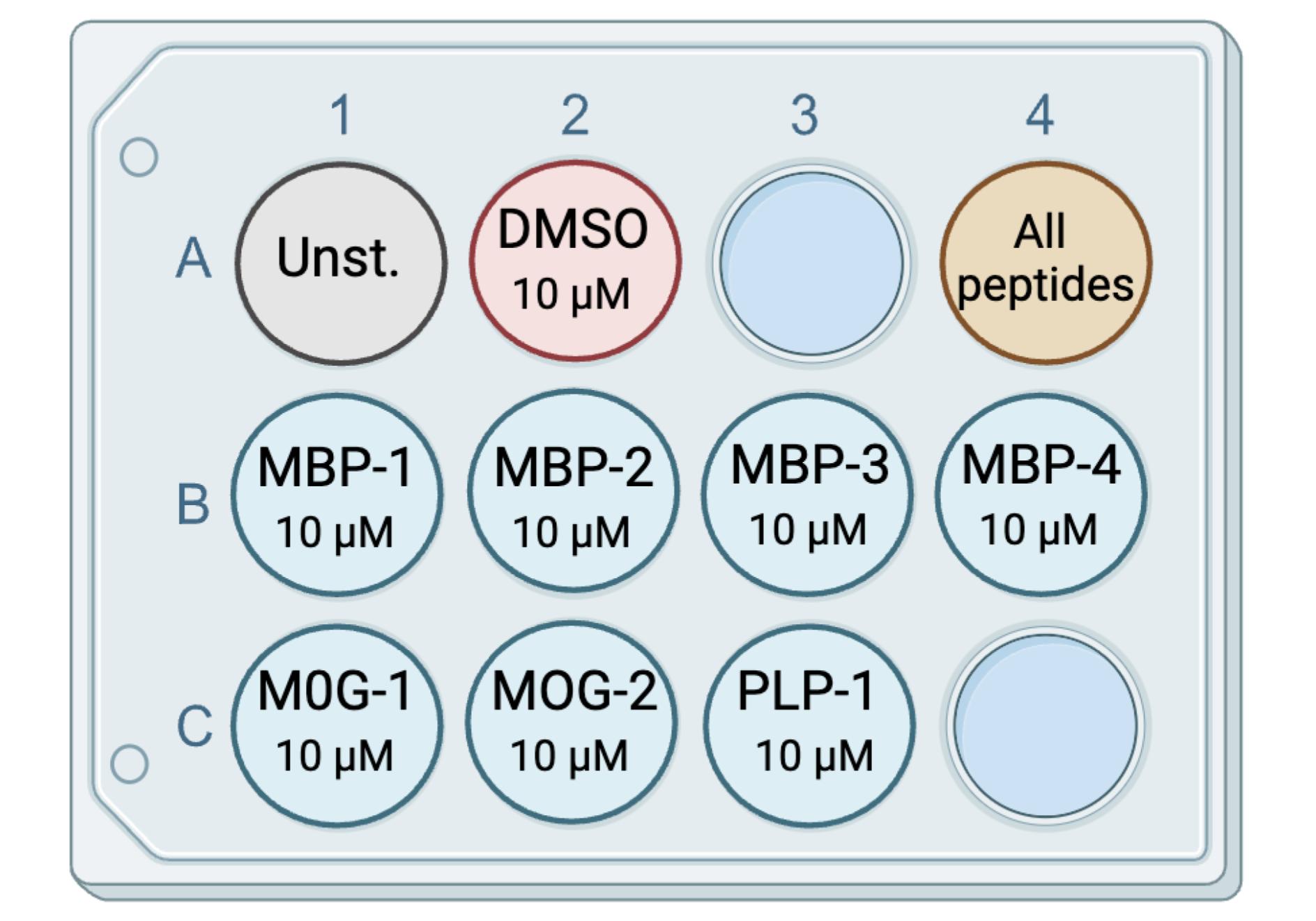

### Supplementary figure 2

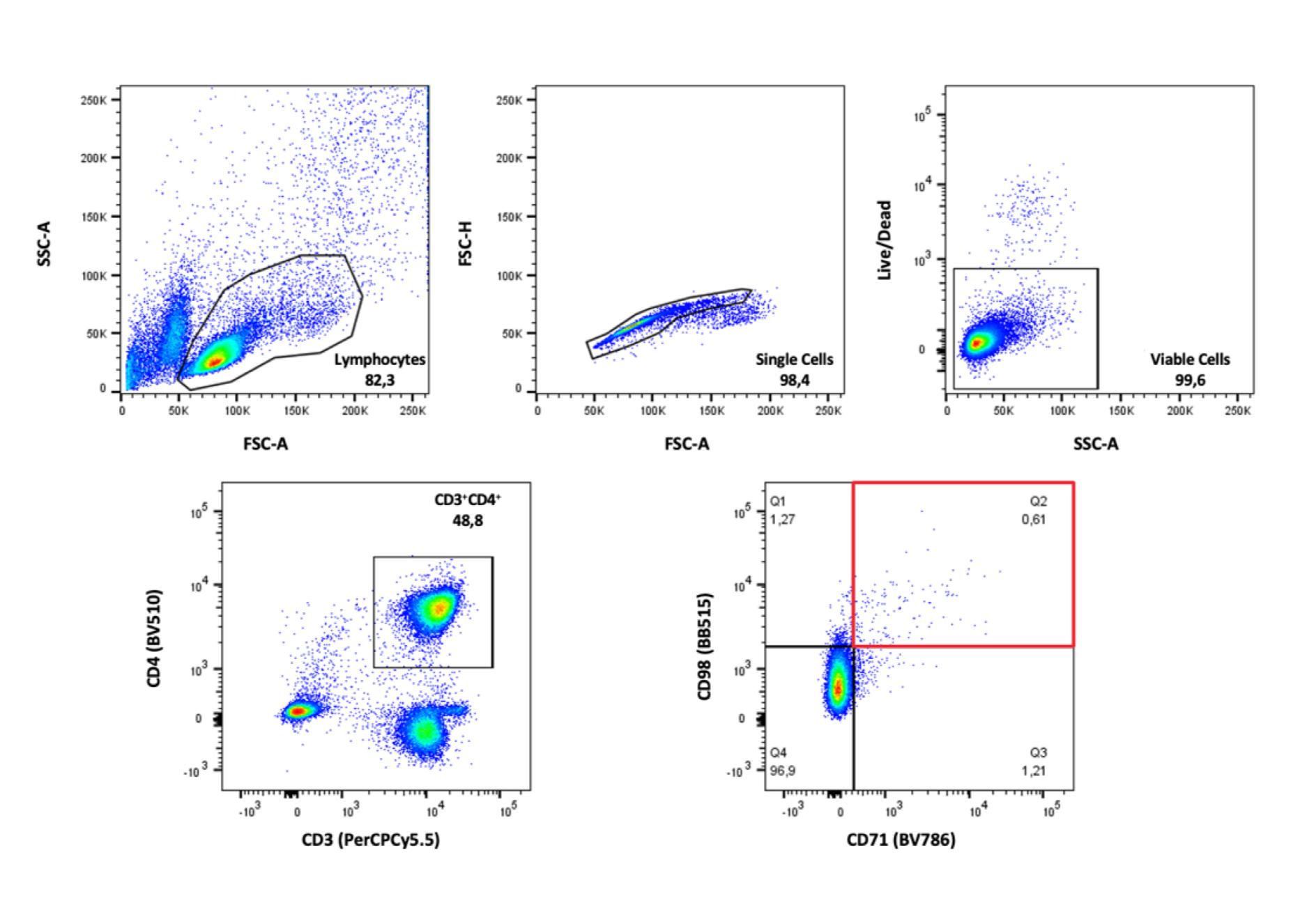

### Supplementary figure 3

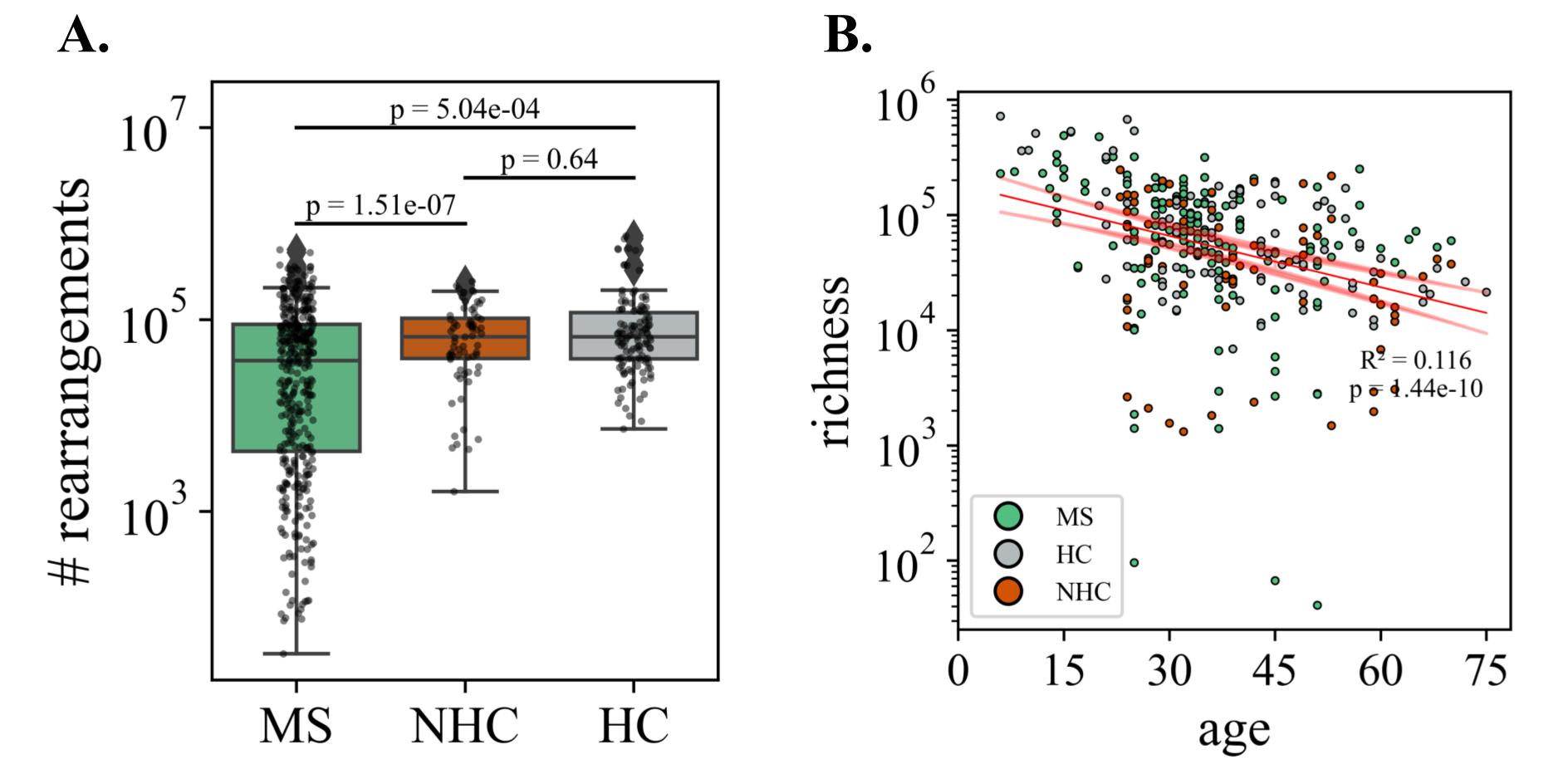

### Supplementary figure 4

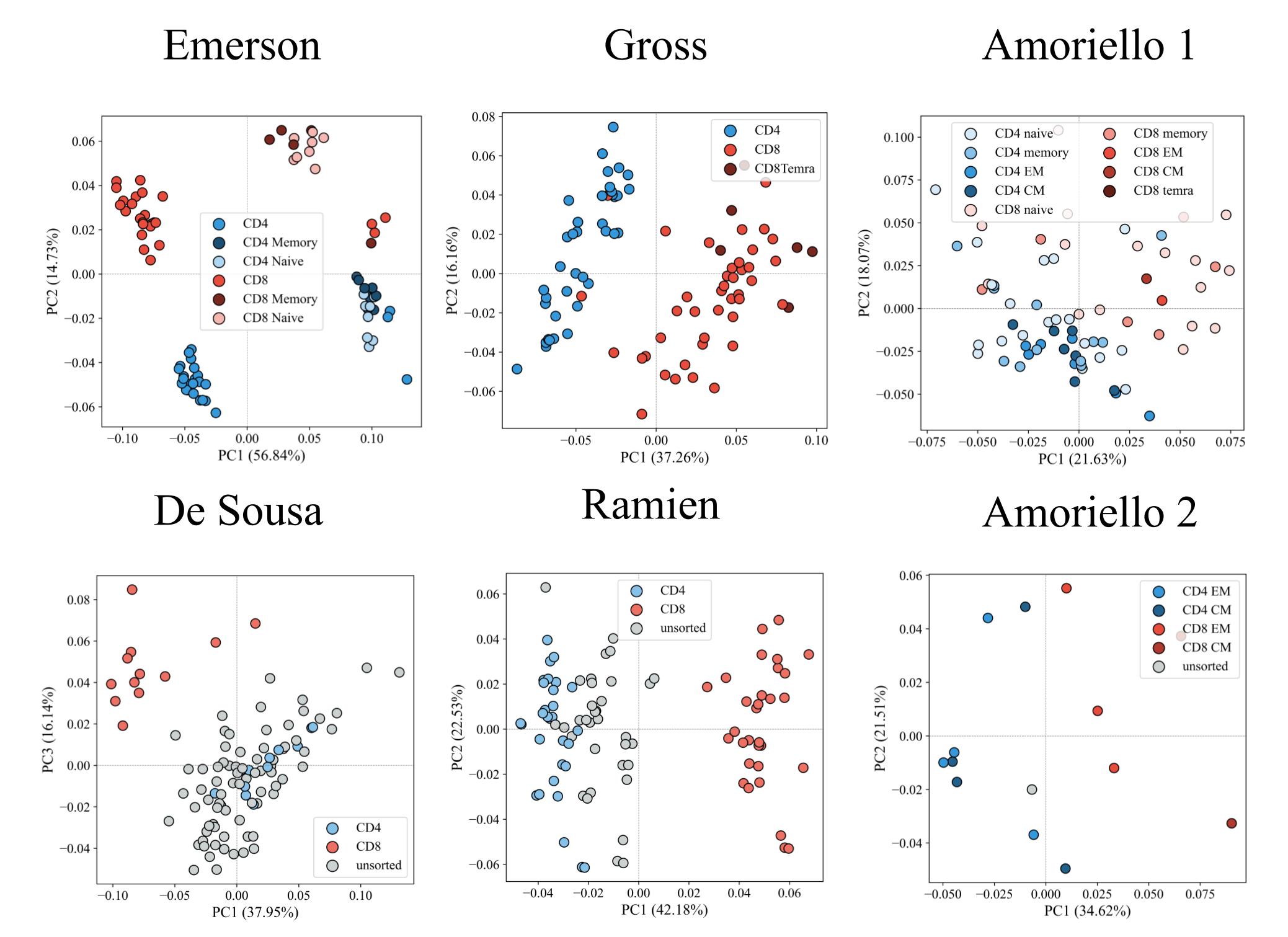

### Supplementary figure 5

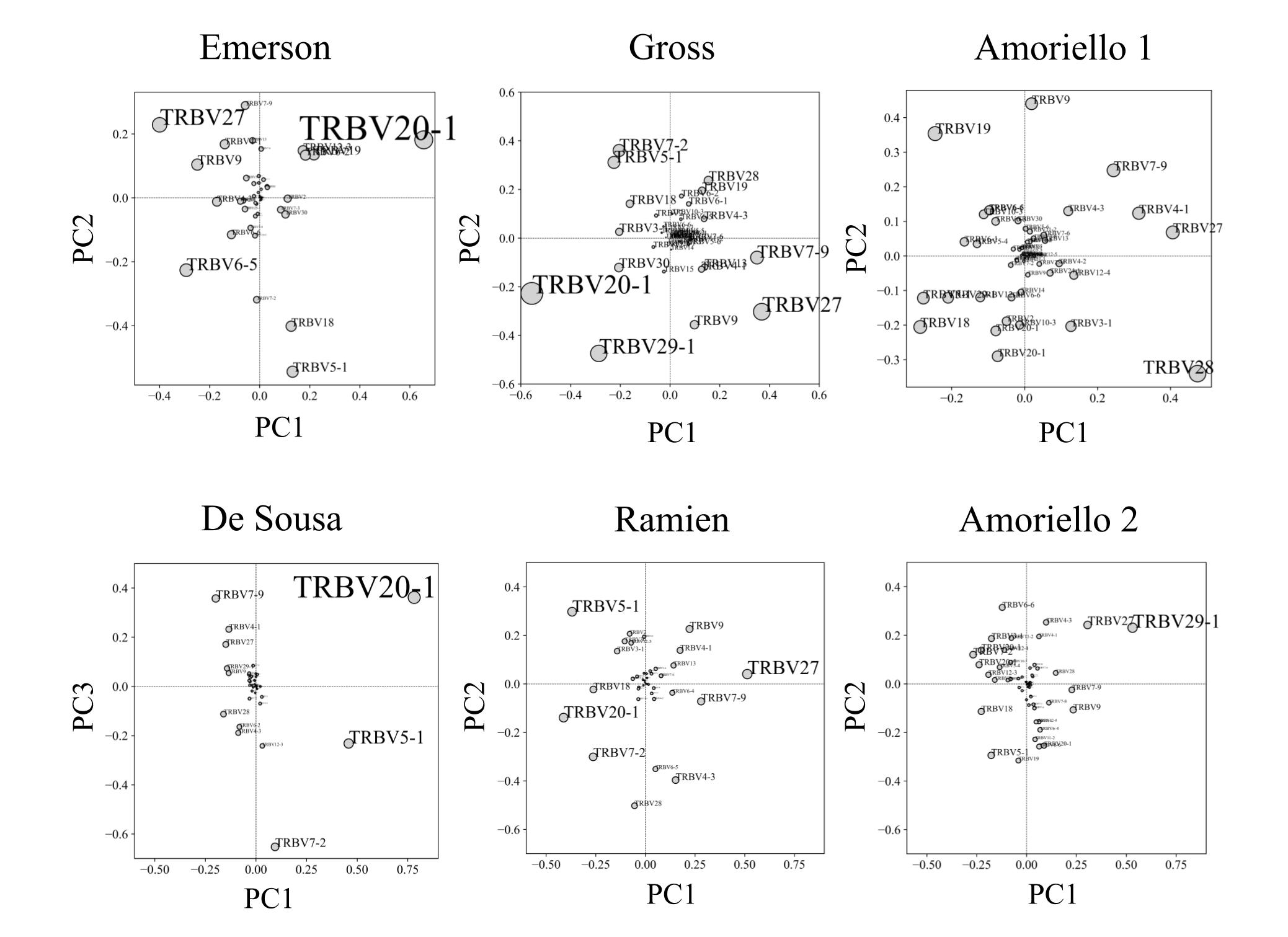

### Supplementary figure 6

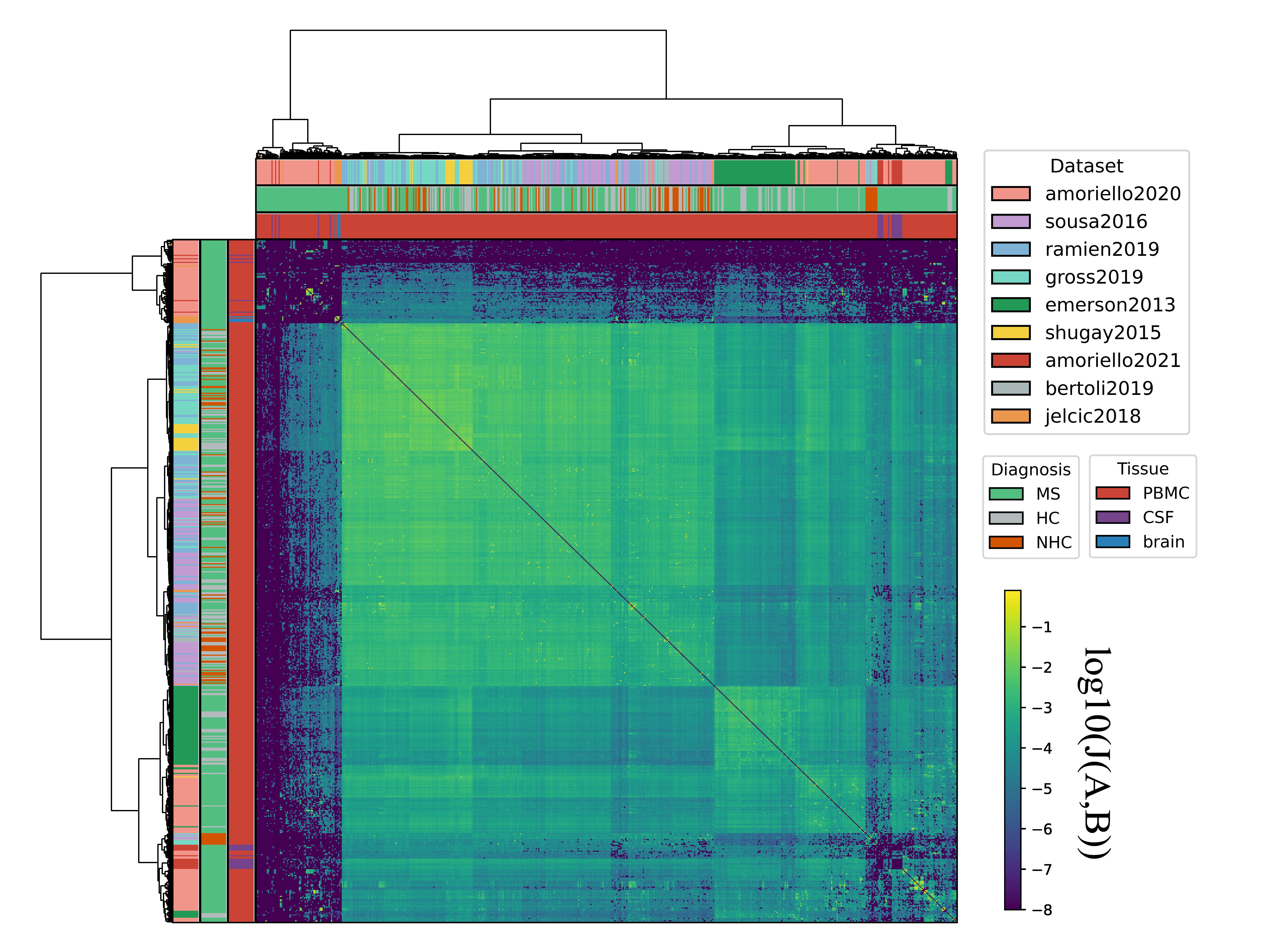

### Supplementary figure 7

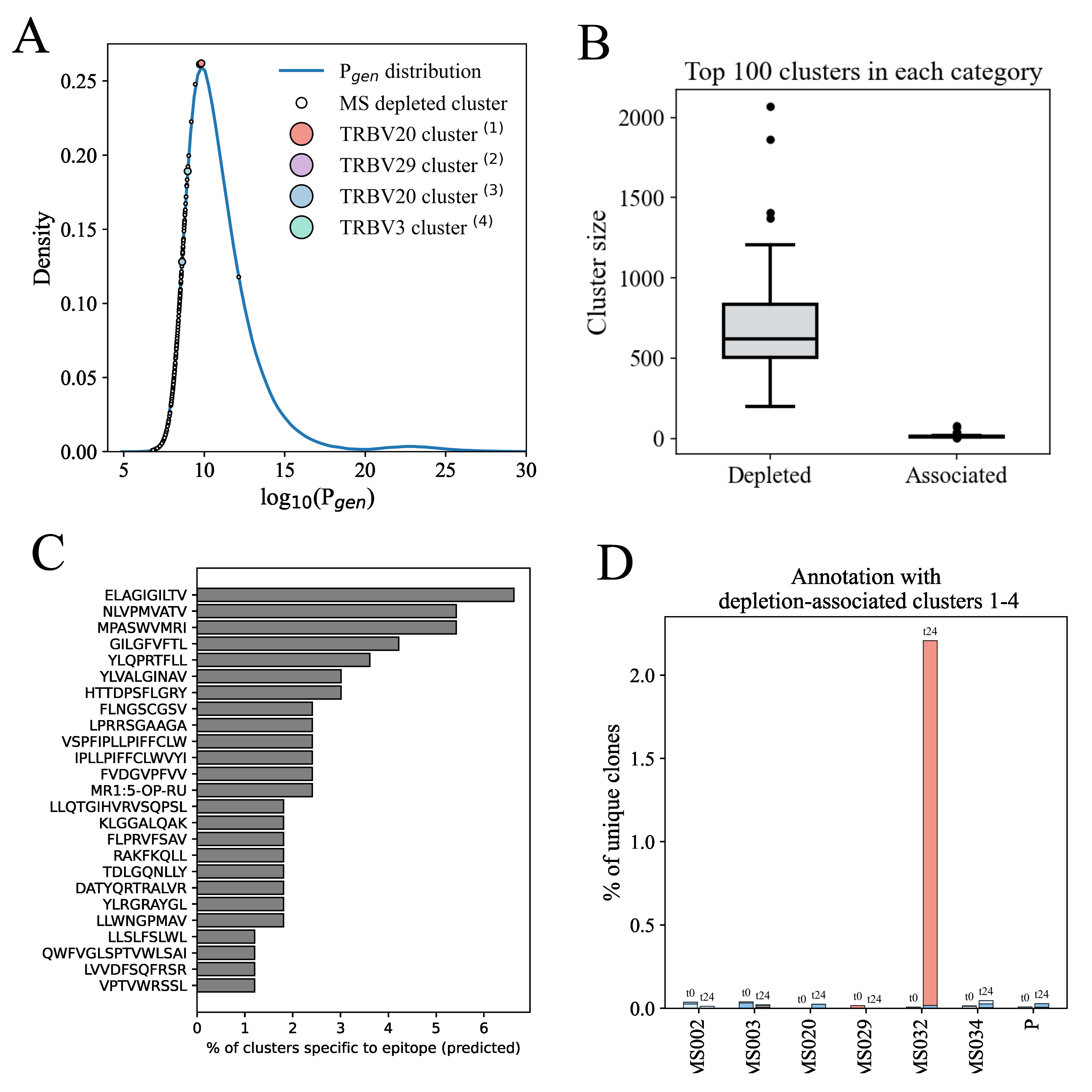

### Supplementary figure 8

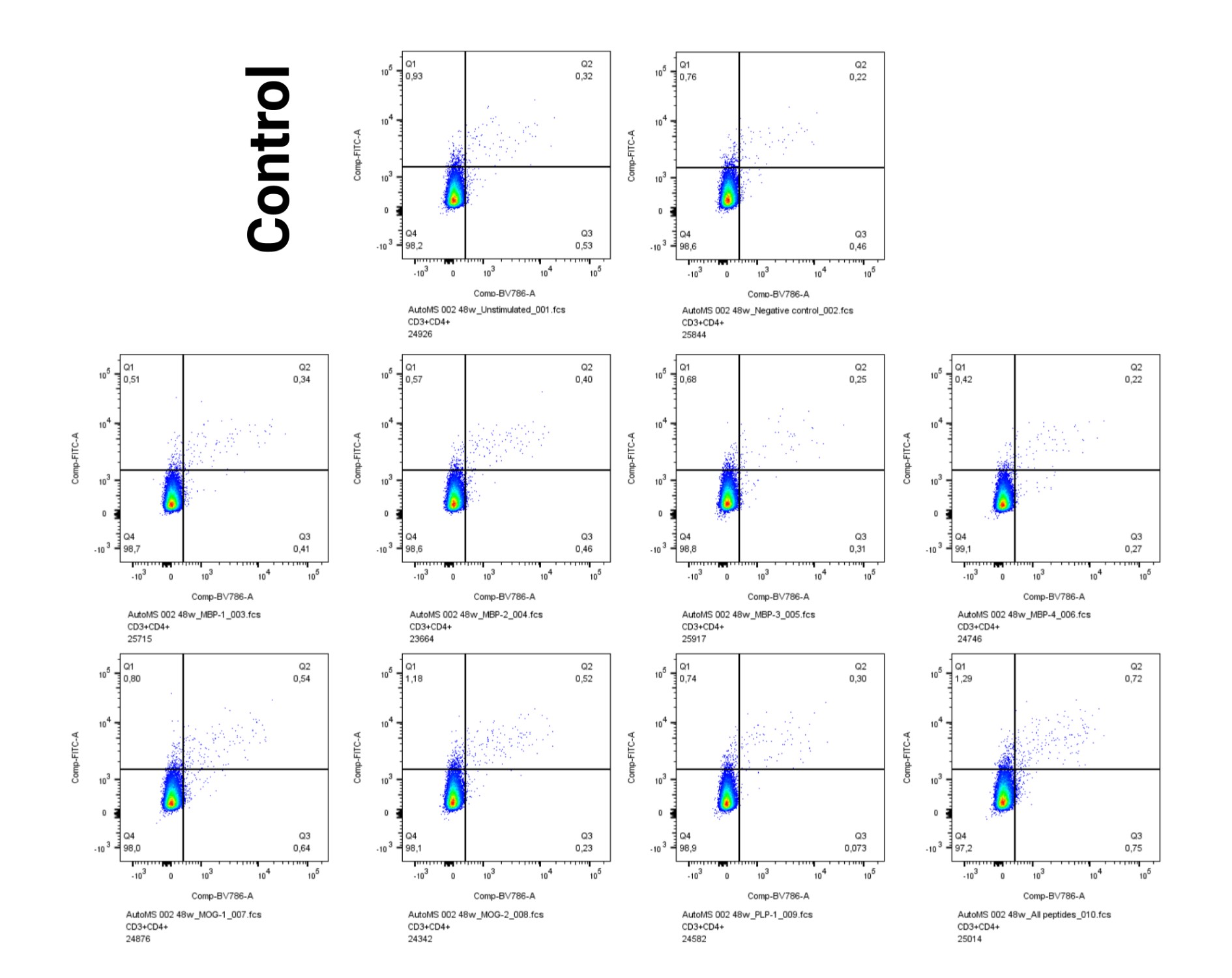

### Supplementary figure 9

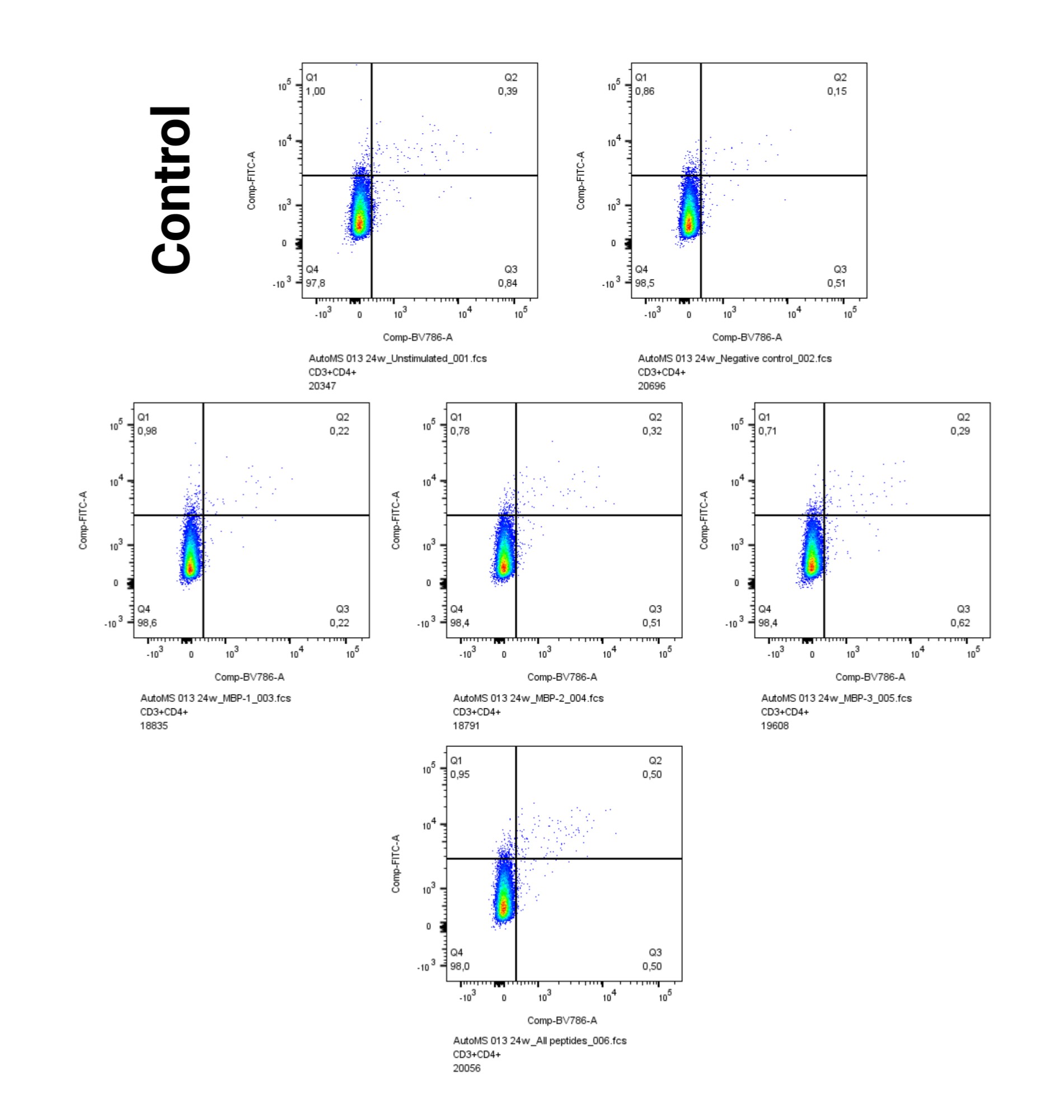

### Supplementary figure 10

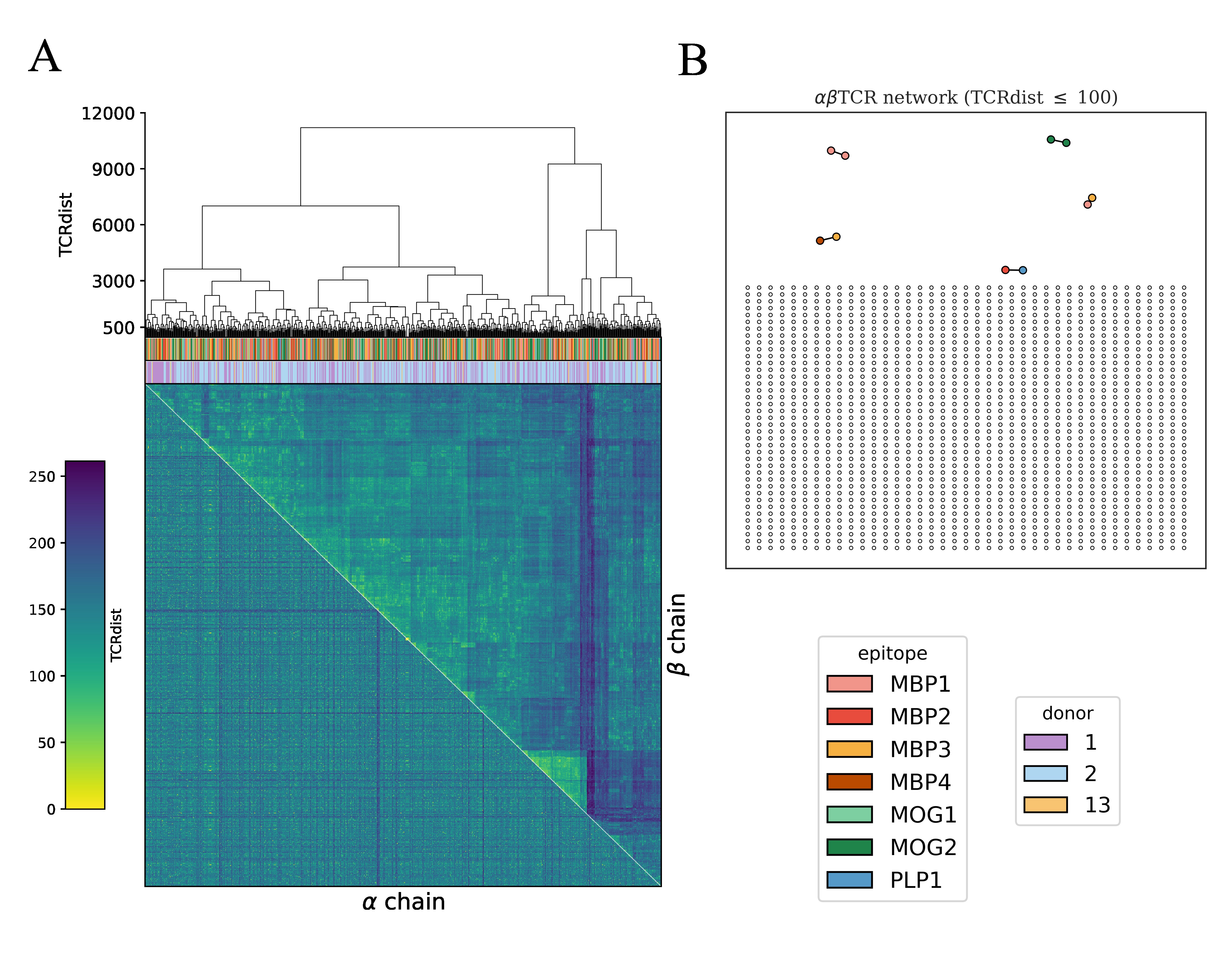

### Supplementary figure 11

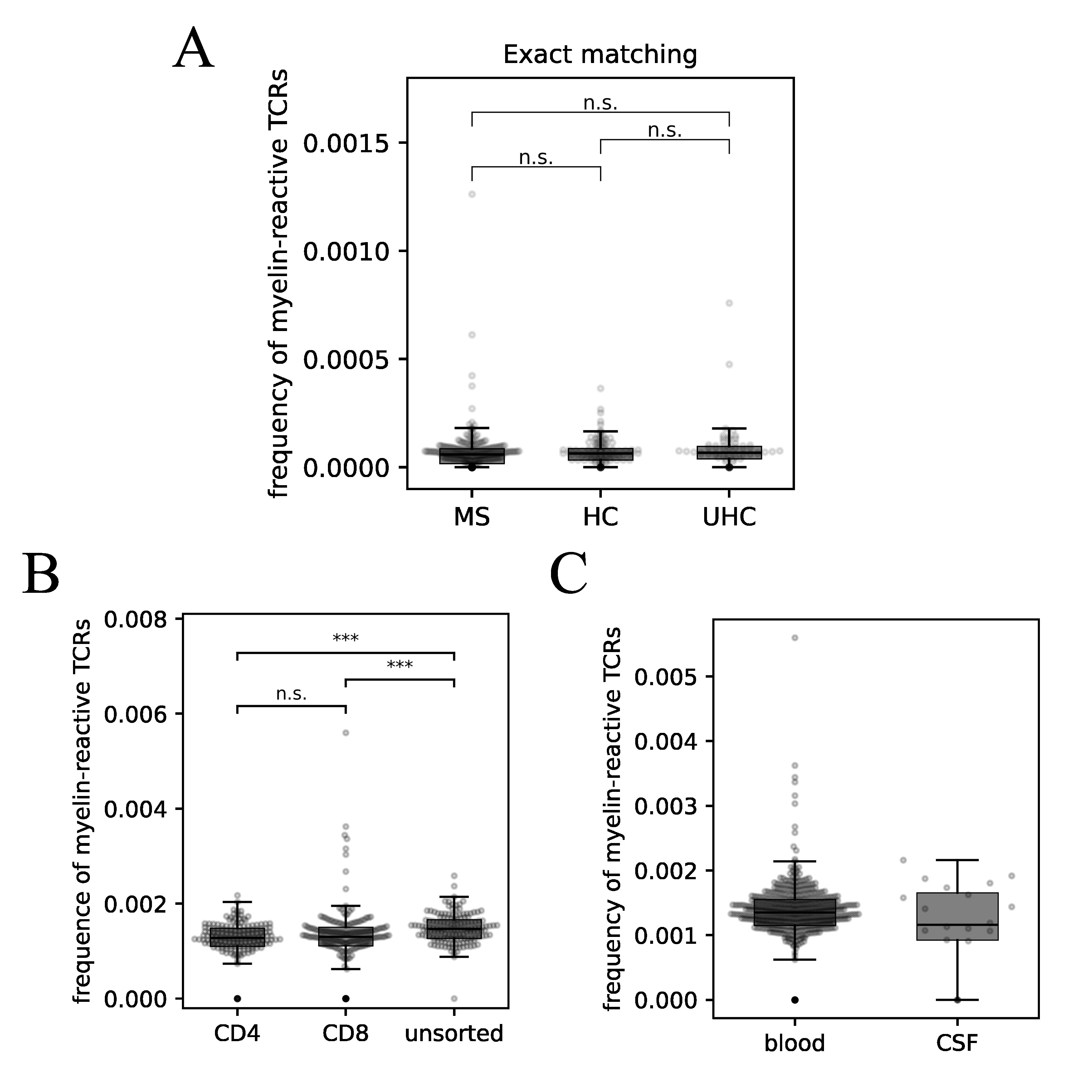

### Supplementary figure 12

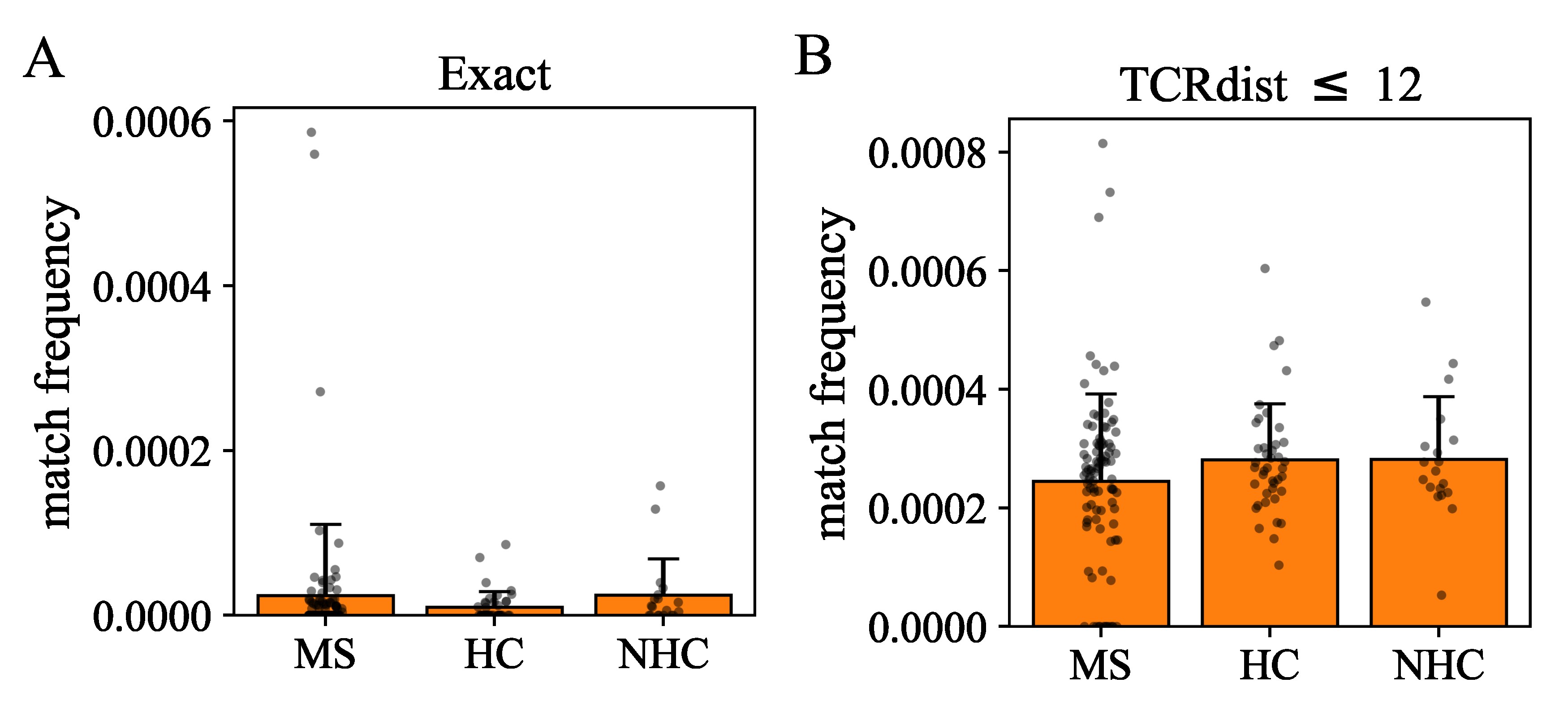
