## Supplementary table 1 for "Linking myelin and Epstein-Barr virus specific immune responses in multiple sclerosis: insights from integrated public T cell receptor repertoires"

| Dataset | Emerson (2013) | Shugay (2015) | Jelcic (2018) | Bertoli (2019) | Gross (2019) | Ramien (2019) | Sousa (2019) | Amoriello (2020) | Amoriello (2021) |
|---|---|---|---|---|---|---|---|---|---|
| Age | No | Yes | No | Yes | Yes | Yes | Yes | No | Yes |
| Sex | No | No | No | Yes | Yes | Yes** | Yes | Yes | Yes |
| Ethnicity | No | No | No | No | No | No | Yes | No | No |
| MS subtype | No | Yes* | No | No | Yes | No | Yes | Yes | Yes |
| Treatment | Unknown | Unknown | Unknown | NTZ, ALEM | Naive | Naive, IFN-β1a, IFN-β1a+ GA, IFN-β1a+ AZA, | Unknown | NTZ, AHSCT | Naive |
| Source | PBMC | PBMC | PBMC, CSF, brain tissue | PBMC | PBMC | PBMC | PBMC | PBMC | PBMC, CSF |
| Cells sorted | Yes | No | Yes | No | Yes | Yes | Yes | Yes | Yes |
| Method | mPCR | 5’ RACE | ImmunoSEQ | ImmunoSEQ | ImmunoSEQ | ImmunoSEQ | 5’ RACE | iRepertoire | iRepertoire |
| Instrument | HiSeq 2500 | HiSeq 2500 | Unknown | Unknown | NextSeq 550 | Unknown | HiSeq 2500 | MiSeq | MiSeq |
| Seq. depth | No | Yes* | No | No | Yes | No | Yes | No | No |
