## Supplementary table 2 for "Linking myelin and Epstein-Barr virus specific immune responses in multiple sclerosis: insights from integrated public T cell receptor repertoires"

| No. | Description oligo-antibody | Manufacturer |
|---|---|---|
| 394661 | TotalSeq$^{TM}$-C0251 anti-human Hashtag 1 | BioLegend |
| 394663 | TotalSeq$^{TM}$-C0252 anti-human Hashtag 2 | BioLegend |
| 394665 | TotalSeq$^{TM}$-C0253 anti-human Hashtag 3 | BioLegend |
| 394667 | TotalSeq$^{TM}$-C0254 anti-human Hashtag 4 | BioLegend |
| 394669 | TotalSeq$^{TM}$-C0255 anti-human Hashtag 5 | BioLegend |
| 394671 | TotalSeq$^{TM}$-C0256 anti-human Hashtag 6 | BioLegend |
| 394673 | TotalSeq$^{TM}$-C0257 anti-human Hashtag 7 | BioLegend |
| 394675 | TotalSeq$^{TM}$-C0258 anti-human Hashtag 8 | BioLegend |
| 394677 | TotalSeq$^{TM}$-C0259 anti-human Hashtag 9 | BioLegend |
| 394679 | TotalSeq$^{TM}$-C0260 anti-human Hashtag 10 | BioLegend |
| 394683 | TotalSeq$^{TM}$-C0262 anti-human Hashtag 12 | BioLegend |
| 394685 | TotalSeq$^{TM}$-C0263 anti-human Hashtag 13 | BioLegend |
| 394687 | TotalSeqTM-C0264 anti-human Hashtag 14 | BioLegend |
| 394689 | TotalSeqTM-C0265 anti-human Hashtag 15 | BioLegend |
