## Supplementary table 3 for "Linking myelin and Epstein-Barr virus specific immune responses in multiple sclerosis: insights from integrated public T cell receptor repertoires"

| **Marker** | **Fluorochrome** | **Clone** | **Reference Number** | **Manufacturer** |
|---|---|---|---|---|
| CD3 | PerCPCy5.5 | UCHT1 | 300430 | BioLegend |
| CD4 | BV510 | RPA-T4 | 300546 | BioLegend |
| CD8 | PB | SK1 | 344718 | BioLegend |
| CD71 | BV786 | M-A712 | 563768 | BioLegend |
| CD98 | BB515 | UM7F8 | 565103 | BioLegend |
| LIVE/DEAD Fixable Near-IR Dead Cell Stain Kit | N/A | N/A | L10119 | ThermoFisher Scientific |
| CD127 (IL-7R) | TotalSeq$^{TM}$-A0390 | A019D5 | 351352 | BioLegend |
| CD25 | TotalSeq$^{TM}$-C0085 | BC96 | 302649 | BioLegend |
