## Supplementary table 5 for "Linking myelin and Epstein-Barr virus specific immune responses in multiple sclerosis: insights from integrated public T cell receptor repertoires"

| patient | timepoint | cluster frequency | cluster count | total TCRs |
|---|---|---|---|---|
| **MS002** | 0 | 0.000086 | 7 | 80995 |
| | 24 | 0.000105 | 13 | 123535 |
| **MS003** | 0 | 0.000194 | 20 | 103035 |
| | 24 | 0.000066 | 4 | 60467 |
| **MS020** | 0 | 0 | 0 | 57889 |
| | 24 | 0.000132 | 4 | 30234 |
| **MS029** | 0 | 0.000031 | 1 | 32247 |
| | 24 | 0 | 0 | 7436 |
| **MS032** | 0 | 0.000038 | 1 | 26351 |
| | 24 | 0.007968 | 381 | 47817 |
| **MS034** | 0 | 0.000071 | 10 | 140925 |
| | 24 | 0.000129 | 14 | 108685 |
| **P** | 0 | 0.000093 | 6 | 64187 |
| | 24 | 0.000161 | 7 | 43588 |
