## Supplementary table and figure captions for "Linking myelin and Epstein-Barr virus specific immune responses in multiple sclerosis: insights from integrated public T cell receptor repertoires"

### Supplementary table legends

**Supplementary table 1.** Detailed overview of public datasets used in this manuscript. NTZ: natalizumab, ALEM: alemtuzumab, IFN-β1a: interferon β1a, GA: glatiramer acetate; AZA: azathioprine, AHSCT: autologous hematopoietic stem cell transplantation. * = information only available for a subset of the participants.

**Supplementary table 2.** Overview of oligo-antibodies. The specificity, reference number and company name are provided for each oligo-antibody.

**Supplementary table 3.** Information of antibodies and dyes used in the flow cytometric experiment used in this study. The specificity, fluorochrome, clone, reference number and company name are provided for each antibody or dye used, when applicable.

**Supplementary table 4**. Linear regression table for age and species richness. The rate of decline in species richness is consistent across the different groups, with species richness decreasing by approximately 1,627 to 5,312 per year of increased age.

**Supplementary table 5.** Presence of selected MS depletion-associated clusters in pre- and post-AHSCT treatment samples. Red indicates an increase in cluster count post-treatment, while blue reflects a decrease

### Supplementary figure legends

**Supplementary figure 1.** Overview of 12-well plate: A minimum of 3.106 PBMCs and maximum of 5.106 PBMCs were co-cultured with nothing (unstimulated), dimethylsulfoxide (DMSO; Sigma-Aldrich, Diegem, Belgium) or stimulated with myelin peptides. (A1) Unstimulated, only PBMCs in this well. (A2) Negative control, PBMCs with DMSO. (A4) All peptides, PBMCs stimulated with all seven myelin peptides with each at a concentration of 10mM. (B1) MBP-1, PBMCs stimulated with MBP-1. (B2) MBP-2, PBMCs stimulated with MBP-2. (B3) MBP-3, PBMCs stimulated with MBP-3. (B4) MBP-4, PBMCs stimulated with MBP-4. (C1) MOG-1, PBMCs stimulated with MOG-1. (C2) MOG-2, PBMCs stimulated with MOG-2. (C3) PLP-1, PBMCs stimulated with PLP-1.

**Supplementary figure 2**. Gating strategy. Gating strategy of CD3+CD4+ cells. First, lymphocytes (SSC-A/FSC-A), single cells (FSH-H/FSC-A) and viable cells (Fixable Near-IR Dead Cell Stain) were selected. After that, T cells (CD3+CD4+) were selected followed by the activated CD4+ cells of interest, with the sort gate indicated in red.

**Supplementary figure 3. A.** Total number of TCR rearrangements for MS, NHC and HC samples. Samples from MS patients contained a lower number of total rearrangements compared to those from NHC (p=1.51e-07) and HC (p=5.04e-04). **B.** Species richness in TCR repertoires of MS, NHC and HC versus age. An age-related decline was observed ($R^2$=0.116, p=1.44e-10).

**Supplementary figure 4.** PCA was conducted on the V gene frequency matrix across the datasets containing T cell subtyping information (six out of nine in total). The combination of the first and second principal components (PC1 and PC2) allows for the visual separation of CD4+ and CD8+ T cell subsets in 2 dimensions in all datasets. For the De Sousa dataset, PC1 and the third principal component (PC3) were used. The axis labels indicate the principal components and the percentage of variance explained by each component.

**Supplementary figure 5.** Loadings plots for V gene usage PCAs. Loadings plots showing the contribution of TRBV genes to the variance in the PCA analysis. The size of each feature (TRBV gene) is proportional to its combined contribution to the first and second principal components (PC1 and PC2).

**Supplementary figure 6. Repertoire similarity of public MS patients' and control subjects' TCR repertoires.** The heatmap represents the pairwise Jaccard similarity between all bulk TCR repertoires, $\log_{10}$-scaled. The columns and rows of the matrix are annotated with dataset of origin, subject diagnosis, and sample tissue type (from top to bottom or left to right, for columns and rows respectively). Rows and columns are linked through hierarchical clustering of the similarity matrix.

**Supplementary figure 7. A.** Pgen of MS-depleted clusters compared to the complete Pgen distribution of the public dataset (blue line). Four selected MS-depleted clusters are indicated with different colours. **B.** Cluster size of top 100 most MS-depleted and MS-associated clusters (by p-value). **C.** Epitope specificity prediction on significantly MS-depleted clusters using IMW DETECT. The x-axis shows the percentage of unique clusters that had predicted specificity (IMW DETECT threshold > 0.23) to the epitope. Top 25 epitopes shown. **D.** Presence of selected MS depletion-associated clusters pre- and post-AHSCT repertoires.

**Supplementary figure 8.** Flow cytometry gating of myelin peptide-reactive TCR pools for donor 013. The first two panels contain 2 control conditions: unstimulated and negative control (DMSO). Flow cytometry gating of myelin peptide-reactive TCR pools for donor 002. The first two panels contain 2 control conditions: unstimulated and negative control (DMSO). The other panels contain the cell gating for MBP1-, MBP2-, MBP3-, MBP4-, MOG1-, MOG2-, and PLP1-activated T cells. The last panel (bottom right) is a combination of all peptides.

**Supplementary figure 9.** Flow cytometry gating of myelin peptide-reactive TCR pools for donor 002. The first two panels contain 2 control conditions: unstimulated and negative control (DMSO). The panels in the second row contain the cell gating for MBP1-, MBP2-, and MBP4-activated T cells. The final panel is a combination of all peptides.

**Supplementary figure 10. Sequence similarity of myelin peptide-activated T cell clones. A.** Annotated heatmap of the pairwise TCRdist distances between the TCRα and TCRβ chains of all myelin-reactive clones. The dendrogram represents the hierarchical clustering based on the TCRβ chain distances. The first row of annotations above the heatmap shows the epitope pool this TCR originated from. The second row represents the different donors. **B.** TCR similarity network of αβTCRs. An edge was drawn between TCRs that had a TCRdist distance of $\leq 100$. Nodes were coloured by myelin epitope. Only nodes with edges are coloured. The legends under the figure were shared for subfigures A. and B.

**Supplementary figure 11.** Annotation of public TCR repertoires from MS patients with myelin-reactive TCRs. A. Comparison of CD4+, CD8+ and unsorted repertoires.

**Supplementary figure 12.** Annotation of CD4+ repertoires from the public database using myelin-specific TCRs from activated CD4+ T cells extracted from subject MS15 (Ramien cohort). The ranksum test showed no statistically significant differences among the different groups.
